# Epigenetic aging of classical monocytes from healthy individuals

**DOI:** 10.1101/2020.05.10.087023

**Authors:** Irina Shchukina, Juhi Bagaitkar, Oleg Shpynov, Ekaterina Loginicheva, Sofia Porter, Denis A. Mogilenko, Erica Wolin, Patrick Collins, German Demidov, Mykyta Artomov, Konstantin Zaitsev, Sviatoslav Sidorov, Christina Camell, Monika Bambouskova, Laura Arthur, Amanda Swain, Alexandra Panteleeva, Aleksei Dievskii, Evgeny Kurbatsky, Petr Tsurinov, Roman Chernyatchik, Vishwa Deep Dixit, Marko Jovanovic, Sheila A. Stewart, Mark J. Daly, Sergey Dmitriev, Eugene M. Oltz, Maxim N. Artyomov

**Author notes:** These authors contributed equally (Irina Shchukina, Juhi Bagaitkar, Oleg Shpynov).

## Abstract

The impact of healthy aging on molecular programming of immune cells is poorly understood. Here, we report comprehensive characterization of healthy aging in human classical monocytes, with a focus on epigenomic, transcriptomic, and proteomic alterations, as well as the corresponding proteomic and metabolomic data for plasma, using healthy cohorts of 20 young and 20 older individuals (~27 and ~64 years old on average). For each individual, we performed eRRBS-based DNA methylation profiling, which allowed us to identify a set of age-associated differentially methylated regions (DMRs) – a novel, cell-type specific signature of aging in DNA methylome. Optimized ultra-low-input ChIP-seq (ULI-ChIP-seq) data acquisition and analysis pipelines applied to 5 chromatin marks for each individual revealed lack of large-scale age-associated changes in chromatin modifications and allowed us to link hypo- and hypermethylated DMRs to distinct chromatin modification patterns. Specifically, hypermethylation events were associated with H3K27me3 in the CpG islands near promoters of lowly-expressed genes, while hypomethylated DMRs were enriched in H3K4me1 marked regions and associated with normal pattern of expression. Furthermore, hypo- and hypermethylated DMRs followed distinct functional and genetic association patterns. Hypomethylation events were associated with age-related increase of expression of the corresponding genes, providing a link between DNA methylation and age-associated transcriptional changes in primary human cells. Furthermore, these locations were also enriched in genetic regions associated by GWAS with asthma, total blood protein, hemoglobin levels and MS. On the other side, acceleration of epigenetic age in HIV and asthma stems only from changes in hypermethylated DMRs but not from hypomethylated loci.

## INTRODUCTION

Advanced age, even in healthy individuals, is accompanied by progressive decline of cognitive, metabolic and physiological abilities, and can enhance susceptibility to neurodegenerative, cardiovascular, and chronic inflammatory diseases^1,2^. Operationally, it is often difficult to determine whether age-associated signatures reflect changes of individual cells or changes in cell type abundances, especially when performing whole tissue transcriptional or epigenetic characterization. For example, T cell numbers are reduced in human blood with age^3^, complicating comparative analyses of whole blood transcriptional or DNA methylation profiles. And even despite a large amount of clinical and epidemiological data^4–8^, we understand very little about the nature of age-associated changes in specific primary cell populations of healthy individuals, particularly with respect to age-associated alterations of the epigenetic landscape. To address this question directly, we focused on classical CD14^+^CD16^−^ monocytes, as they are homogeneous, easily accessible, and relatively abundant in blood, which permits multi-omics profiling of these cells obtained from a single blood draw. Also, monocytes have a short differentiation trajectory from bone marrow progenitors, rendering them informative proxies to understand epigenetic changes that occur with age in human hematopoietic stem cells.

Epigenetic aging can manifest in two key aspects: via age-associated changes in chromatin modifications and in DNA methylation. Robustness of the connection between aging and DNA methylation has been well acknowledged^9–13^, yet in spite the large number of studies, cell specific regions of age-associated DNA methylation/demethylation have not been reported to date. Previous studies have been predominantly using DNA methylation arrays which detect changes of a predefined set of distant solitary cytosines across the genome^5^. This design, however, inherently prevents identification of differentially methylated regions (DMRs), which are expected to be more biologically relevant compared to changes in single isolated CpG sites. Furthermore, while total level of various chromatin modifications has been characterized across age groups and cell populations using CyTOF^14^, no genomically resolved profiling of major chromatin modifications (e.g. via ChIP-seq) has been reported for primary human cell types and extent of chromatin changes in primary human cells during aging remains unclear.

In this study, we used parallel multi-omics approaches to characterize intracellular states and extracellular environments of monocytes along healthy aging. We profiled DNA methylation, five major chromatin modifications, as well as transcriptional and proteomic states of purified CD14^+^CD16^−^ monocytes from 20 young (24-30 y. o.) and 20 old (57-70 y. o.) donors. To allow identification of continuous DNA regions that change methylation level with age and comprehensive characterization chromatin state, we utilized enhanced Reduced Representation Bisulfite Sequencing (eRRBS) coupled with the Ultra-Low-Input ChIP-Sequencing (ULI-ChIP-seq)^15^ approach to profile chromatin modifications from limited input material. Our approach led to identification of more than a thousand DMRs, which could not be achieved via methylation arrays technology (only ~3% of the DMRs are sufficiently covered in the most commonly used methylation array design). We found no evidence of large-scale remodeling of chromatin modification landscape along the healthy aging yet revealed distinct chromatin features that were characteristic of age-associated DNA hyper- and hypomethylated regions. Integration of the obtained DMR signatures with transcriptional data highlighted connection between age-associated transcriptional alterations and hypomethylated DMRs, while hypermethylated DMRs were not readily associated with transcriptional changes. Together with parallel profiling of plasma proteins and unbiased metabolic profiling from the same individuals, the compendium of data collected in this work comprises a comprehensive aging data resource obtained under stringent inclusion criteria. Easily accessible visualization and exploration of all data from this study, and detailed descriptions of protocols and computational pipelines are available online at http://artyomovlab.wustl.edu/aging/.

## RESULTS

### Study design, cohort characteristics, and systemic age-associated changes

It has been described that genetic factors, race, sex, body-mass and lifestyles can significantly affect the outcomes of human studies focused on various aspects of aging^7,8,10,16,17^. Here, we have attempted to assemble a cohort as thoroughly controlled as possible to limit non-age-related contributions in the downstream characterization of monocytes. We used stringent inclusion criteria to eliminate the effect of confounding variables such as inherited genetic traits, sex, environmental stressors and inflammatory disorders^18,19^. Blood was collected from healthy young (24-30 y.o.; n=20) and old (57-70 y. o.; n=20) Caucasian males. All donors were non-smokers, with a healthy body-mass ratio (BMI<30) and self-reported absence of underlying inflammatory conditions, acute viral infections, and cancer (Figure 1A, **Table S1**). Since we focused on profiling of immune cells, blood pressure medication was allowed by inclusion criteria in the older cohort. We additionally validated non-inflamed status of all recruited donors by profiling pro-inflammatory cytokines in plasma using a bioplex assay (**Figure S1A; Table S2**). All cytokines were within normal ranges^20^, confirming that our selection process was sufficiently stringent and excluded underlying inflammatory conditions in both young and old participants^21–23^. One cytokine that stood out was IL-8 that was strongly statistically increased with age even for the considered cohort size (while still remaining within the healthy range, **Figure S1A**).

**Figure 1:**
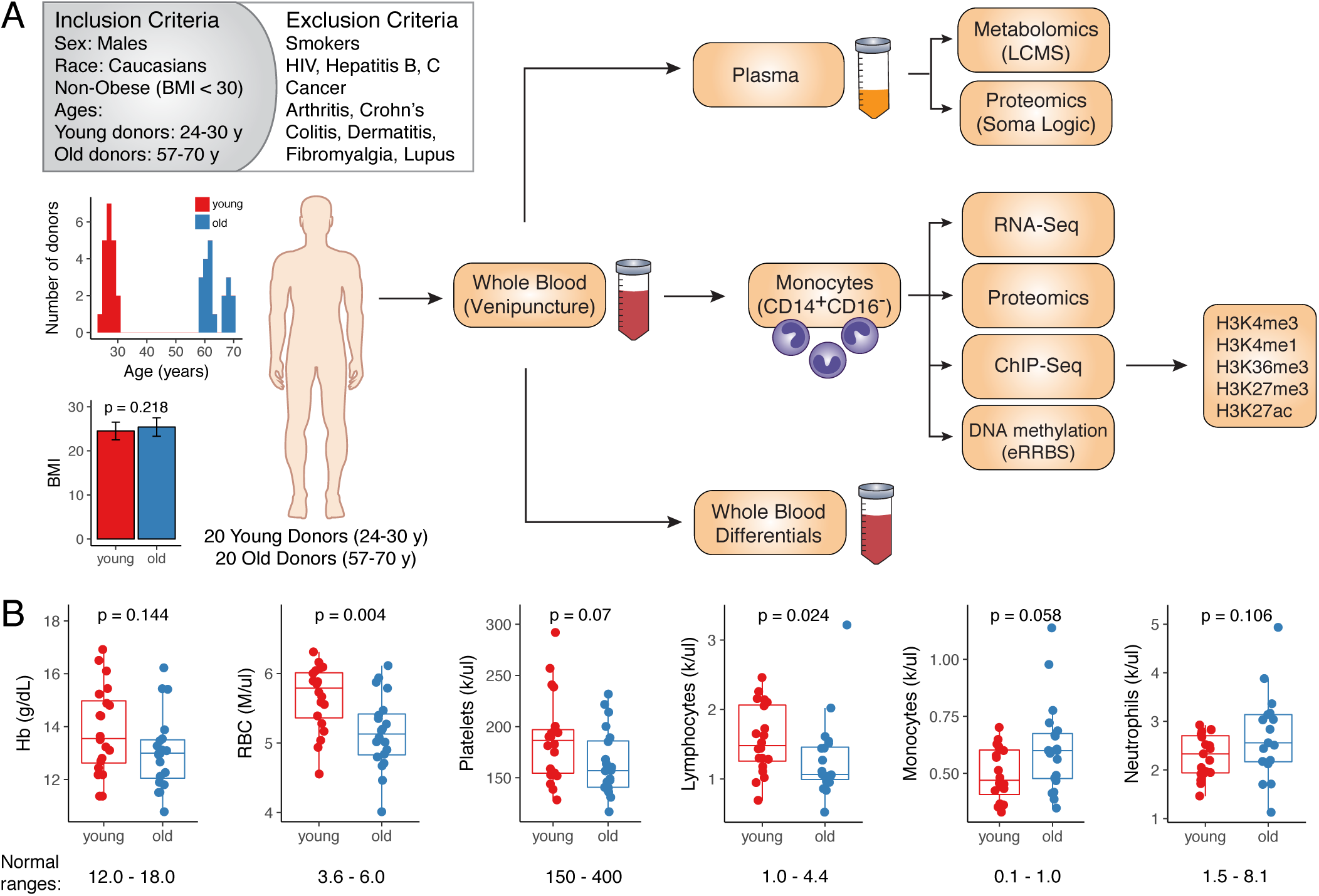
Integrated multiomics profiling of healthy aging: study design overview. **(A)** Overview of selection criteria and study design. Histogram represents distribution of ages amongst young (n=20, red) and old (n=20, blue) donors. BMIs of donors are show as mean ± S.D. P-value was calculated using two-sided Mann-Whitney U test. **(B)** Blood differentials obtained by Hemavet. Normal ranges are shown below the boxplots. Statistical differences between groups were estimated using two-sided Mann-Whitney U test

Complete blood counts (CBCs) and differential analysis of whole blood (Figures 1B **and S1B; Table S1**) showed a significant decrease in red blood cell (RBCs) counts with age, consistent with previous reports^24^. This decrease was not categorized as anaemia, since haemoglobin and haematocrit levels were within a healthy range (Figures 1B and **S1B**). Total white blood cell (WBC) counts were similar between the young and old cohorts but we saw significant changes in WBC differentials: total lymphocyte counts decreased with age, while myeloid cell counts were higher, with monocytes being more abundant in the older cohort (Figure 1B and **S1B**). These data are consistent with previous reports of age associated shifts towards the myeloid lineage^25^.

To characterize the differences in the environment that bathes circulating monocytes, we performed proteomic and metabolomic profiling of plasma (**Figures S2 and S3**). PCA and hierarchical clustering revealed moderate separation between the two cohorts for both datasets (**Figures S2A, 2B and S3A, 3B**). Statistical testing resulted in 39 significantly different metabolites (FDR < 0.05; **Figure S2C; Table S3**) and 53 significantly different proteins (FDR < 0.05; **Figure S3C; Table S4**). Overall, age-associated protein signature corroborated a recently published reports by the Ferrucci group^26,27^, e.g. sclerostin (SOST) and growth differentiation factor 15 (GDF15) were among the most distinct proteins, and we additionally validated them and three other targets by ELISA (**Figure S3D; Table S5**). Metabolomic data analysis showed a dramatic decrease in sex steroid levels for the older population, consistent with previous publications^28^ (**Figures S2D and S2E**). Altogether, pathway analysis identified statistically significant age-associated changes in 15 metabolic pathways (**Figures S2F and S2G**). Interestingly, apart from the expected steroid-related and secondary bile acid pathways^23,29^, phospholipid metabolism pathway was also downregulated with age. Eleven pathways that were upregulated in the older cohort include pathways involved in metabolism of xanthine and caffeine, benzoate, nucleotides, peptides, assorted amino acids, fatty acids, and non-glycolytic carbohydrates. Finally, for correlation analysis between proteomic and metabolomics datasets we focused on age-associated proteins and discovered that some plasma metabolites were co-regulated with protein markers only in the old group (**Figure S3E**). For instance, GDF15 and NOTCH1, both strongly associated with age, were highly correlated with cystine and glucuronate in the old cohort but not in the young (**Figure S3F**). Overall, the plasma profiling data validated our recruitment criteria as well as confirmed and expanded previously known systemic features of aging. We next aimed to understand intracellular determinants of aging by focusing on CD14^+^CD16^−^ monocytes.

### Protein levels change substantially while transcripts levels are more robust during healthy aging

To explore intracellular signatures, we profiled pure CD14^+^CD16^−^ monocytes (>98% purity; **Methods**, **Figure S4A**) from young and old individuals using deep RNA-sequencing (**Table S6**). PCA of these data showed no clear separation of old and young transcriptomes and differential expression analysis revealed few significantly changing genes (Figures 2A and 2B, **Table S7**). One possible reason for this could be that cohort sizes in our study were limiting the power to detect changes of small absolute magnitude. Thus, we re-analyzed a large publicly available dataset generated as a part of the MESA study, which profiled purified monocyte samples from over 1,200 donors between the ages of 44 and 83 years^30,31^. Consistent with the initial report^31^, we identified 4,549 statistically significant differentially expressed genes (Figure 2C; **Table S8A**). Yet, fold changes of gene expression between middle-aged and old MESA donors were very small – most changes being around 1% of the average expression level of the corresponding gene (Figure 2D). Our down-sampling simulation showed that to detect changes of this magnitude one requires a dataset with at least ~100 donors per group (Figure S4B). Therefore, we conclude that both our and public data demonstrated the relative stability of transcriptional landscape in human monocytes characterized by small magnitude age-associated changes.

**Figure 2:**
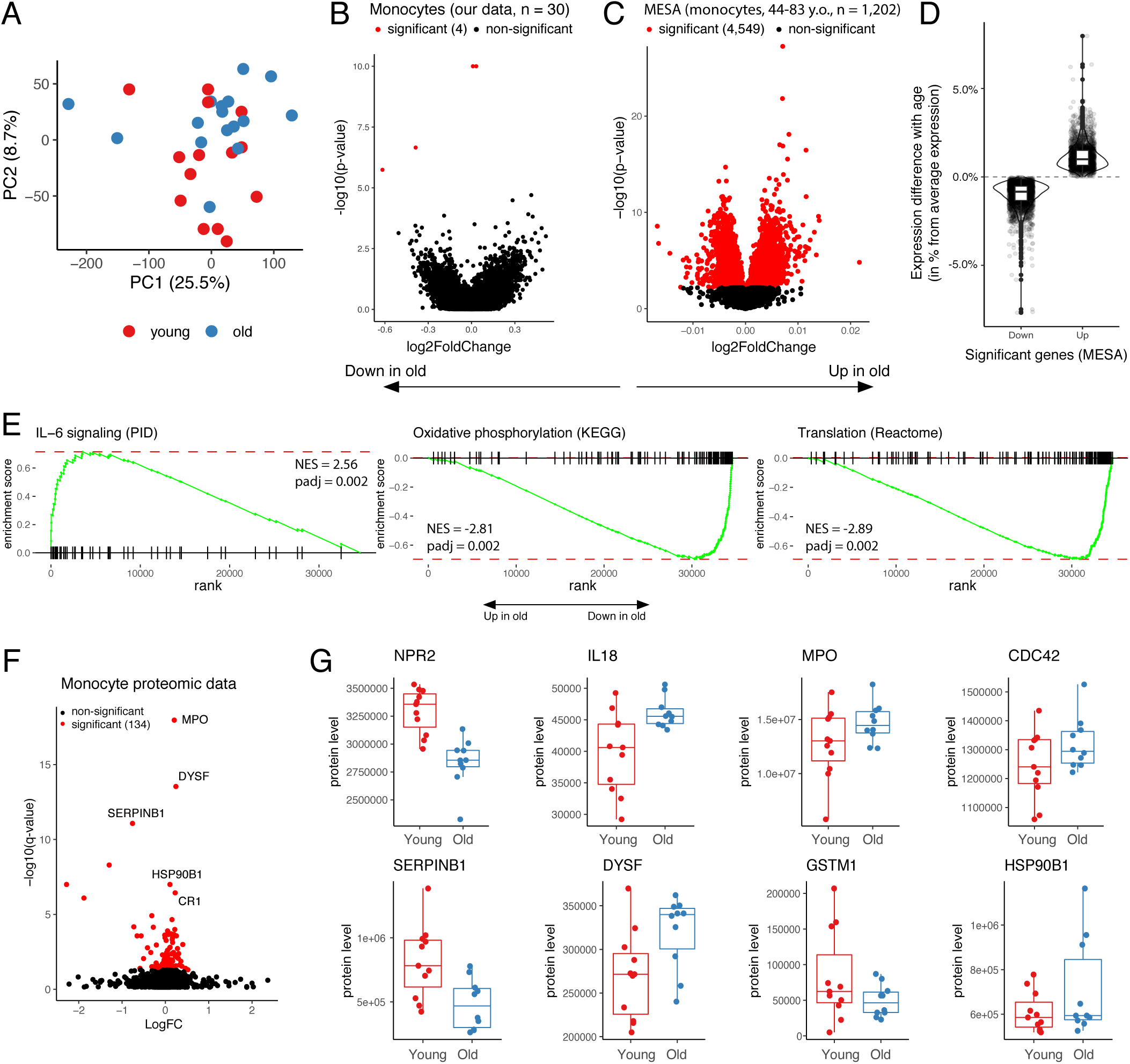
Healthy aging in monocytes: very low magnitude transcriptional changes and distinct proteome alterations. **(A)** PCA of normalized expression levels estimated by RNA-seq. Each dot represents one donor, percentage of variance explained by principal components is shown in brackets. **(B)** Volcano plot depicts results of differential analysis. Each dot represents a single gene. P-values and logFC were calculated by DeSeq2 package. Significant genes are highlighted in red. **(C)** Volcano plot as in (B) represents differential expression results for MESA transcriptomic data for 1,202 human monocyte samples (GSE56045). P-values and logFC calculated by Limma package. **(D)** Difference in mean expression between the most extreme MESA age groups (top and bottom 10% quantile) was presented as percent from average expression level of the corresponding gene across all donors. Only significant genes were included. **(E)** GSEA enrichment curves illustrate examples of pathways significantly up- and down-regulated with advancing age in MESA data. **(F)** Volcano plot shows result of monocyte proteome differential analysis. Each dot represents a protein. **(G)** Examples of age-dependent monocyte proteins.

In the case of small absolute changes, gene set enrichment analysis (GSEA) is a useful approach to find coherently changing pathways. Indeed, we identified a number of significantly altered with age pathways (**Table S8B**), including increase in cytokine signaling pathways, a decrease in oxidative phosphorylation pathway, and decrease in multiple protein translation pathways (Figures 2E and **S4C; Table S8B**). The latter led us to an idea that more profound changes between age groups might be present at the level of proteome. Therefore, we subjected monocytes to proteomic profiling, which detected significant age-associated changes in 134 proteins (Figures 2H and 2I; **Table S9**). We found an increase in protein levels of natriuretic peptide receptor (NPR2), cytokine interleukin 18 (IL-18), myeloperoxidase (MPO), and toll-like receptor chaperon heat shock protein 90 beta family member 1 (HSP90B1) in the older cohort, suggesting that baseline condition of monocytes might be skewed towards more pro-inflammatory state in older individuals. We also report an increase in protein level of CDC42 in older cohort. This dysregulation might stem from the HSC level, since CDC42 inhibition has been reported to functionally rejuvenate aged HSCs in mice^32^. Interestingly, both IL-18 and GSTM1 were previously associated with age-related phenotypes in genome association studies^33,34^. Analysis of age-associated changes in gene expression corresponding to the significant proteins revealed that majority of the identified protein level alterations could not be explained by a shift in expression, suggesting age-associated disturbance of post-transcriptional regulation (**Figure S4F**).

The difference in monocytes’ proteome suggested that despite similarity of their transcriptional profiles old and young monocytes might respond differently to activating stimuli. We tested this hypothesis using lipopolysaccharide (LPS) stimulation of monocytes from young (n=7) and old donors (n=7) and macrophages *in vitro* differentiated from these monocytes. We collected and sequenced RNA from these four groups of samples (52 samples in total) (**Figure S2D, Table S10**). While we observed known transcriptional signatures of monocyte differentiation and activation, we again could not detect any significant changes between activated cells from different age groups (**Figure S2E**). Similar to baseline monocyte profiling, this does not necessarily imply the absence of the age-associated changes but indicates that their absolute magnitude is likely low and requires bigger cohorts to be detected. Additionally, larger changes might still be observed in the case of monocyte differentiation into other lineages, such as dendritic cells and/or osteoclasts, or with usage of different activating stimuli. *Taken together, our data show that transcriptional profiles of monocytes do not change considerably along the healthy aging, while larger magnitudes of changes are observed on the protein level.*

### Identification of differentially methylated regions (DMRs) associated with healthy aging

A number of studies establish robust relationship between age and DNA methylation using DNA methylation arrays in various cell types or whole tissues^9,10,12^. These observations were underscored by the development of DNA methylation-based algorithms for age prediction^11,35–40^. Yet, DNA methylation array technology is limited in ability to find dedicated regions undergoing age-related DNA methylation or demethylation. Here, we used enhanced reduced representation bisulfite sequencing (eRRBS)^41^, which allows for identification of continuous regions by sequencing many closely located cytosines, providing deeper insight into the DNA methylation landscape. Individual eRRBS libraries for each sample were sequenced at 70±10 million reads depth (**Figure S5A, Table S11**) and yielded approximately 3 million well-covered CpGs (with mean coverage ≥10 reads across all samples). This coverage represents ~10% of all CpGs in the human genome, including 24,127 CpG islands (84% of all islands, Figure 3A).

**Figure 3:**
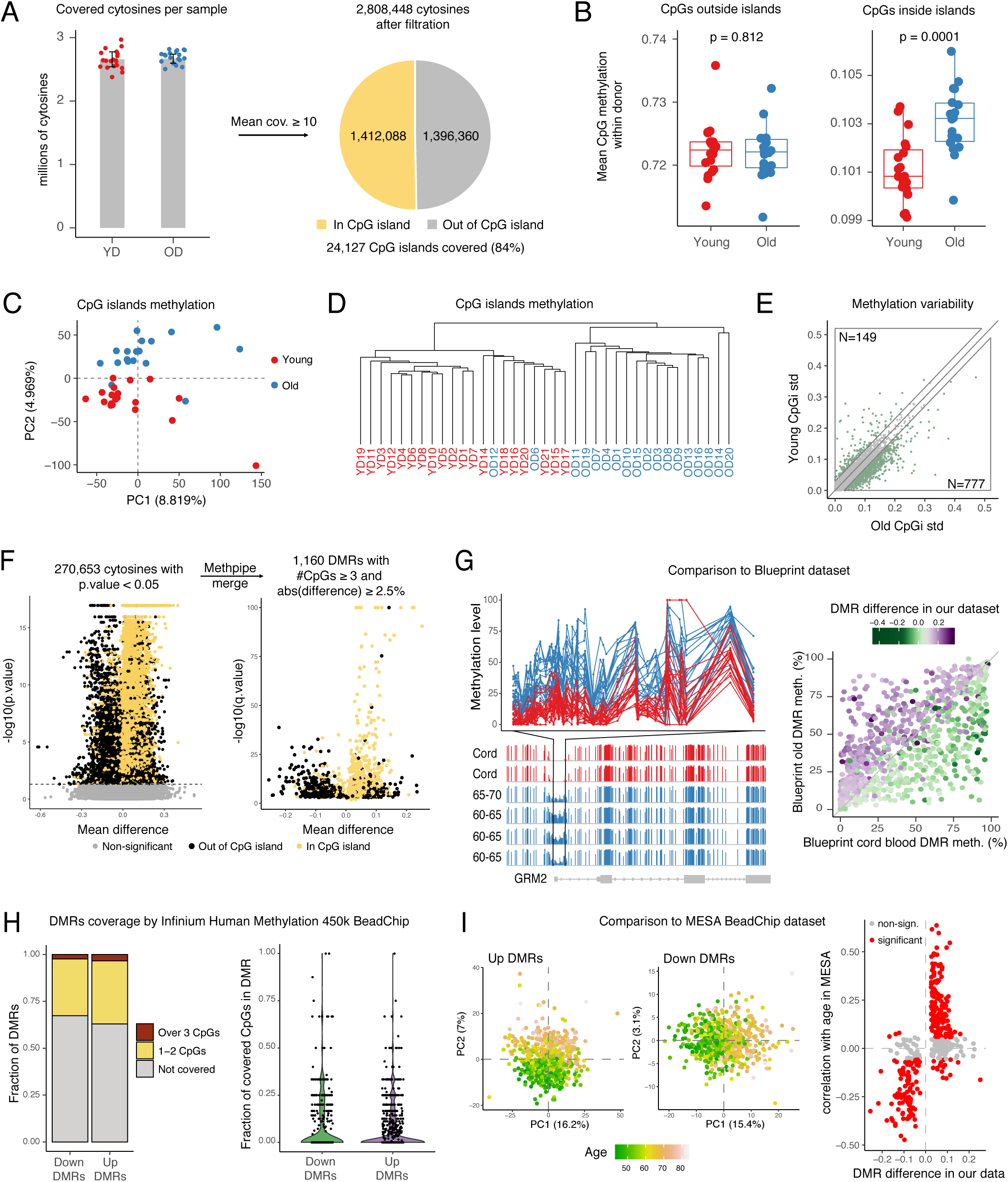
RRBS sequencing reveals robust continuous regions of age-associated DNA hyper- and hypomethylation (up and down DMRs). **(A)** Number of covered cytosines across both cohorts. Approximately 2.8 million cytosines remain in analysis after filtration (average coverage ≥ 10 reads), covering most CpG islands. **(B)** Comparison of global methylation levels between age groups. Each dot represents mean methylation of all cytosines outside (left) or inside (right) CpG islands in one sample. P-values were determined by 2-way ANOVA. **(C)** PCA of scaled CpG island methylation levels. Each dot represents one sample, percentage of variance explained by principal components is shown in brackets. **(D)** Dendrogram produced by unsupervised hierarchical clustering of all samples. Each sample described as a vector of CpG island methylation levels. Clustering using Ward algorithm and Manhattan distance as distance metric. **(E)** The scatter plot represents the standard deviation (std) of every CpG island within old (x-axis) and young (y-axis) cohorts respectively. Upper (top left) and lower (bottom right) areas highlight islands having higher variability in young and old cohorts respectively. Numbers of both are shown. **(F)** Differentially methylated region (DMR) detection. Each dot on the left volcano plot represents one cytosine. P-values and methylation differences (average(old)-average(young)) were calculated using MethPipe toolkit. Yellow color indicates that cytosine is located within a CpG island. Cytosines with p-values over 0.05 considered non-significant (grey dots). Consecutive significant cytosines were merged into DMRs using Methpipe (right volcano plot). DMR p-values were calculated by Fisher method. Yellow dots represent DMRs that intersect CpG island. **(G)** Comparison to Blueprint dataset. Left: example of DMR in GRM2 promoter. Upper panel shows % methylation level (in percent, y axis) of each cytosine inside the DMR defined by our data (x axis represents position in genome). Lines represent young (red) and old (blue) donors. Bottom panel shows Blueprint WGBS tracks in the corresponding region for classical monocytes from cord blood (n=2, red) and old donors (n=4, 60-70 y. o., blue); y axis represents cytosine methylation levels. Right panel: each dot represents one DMR detected in our dataset. Dots above diagonal gain methylation with age according to the Blueprint dataset, dots below – lose it. Color shows the difference in region methylation between young and old groups in our dataset (green – hypomethylation, purple – hypermethylation). **(H)** Summary of DMR coverage in methylation array. On the left panel each dot represents a single DMR **(I)** Comparison to MESA dataset. Left: PCA of up and down DMR methylation levels in MESA dataset. Each dot represents one sample, percentage of variance explained by principal components is shown in brackets. Dots are colored according to donor age. Right: Spearman correlation of DMR methylation and age was calculate for all DMRs using MESA dataset. Each dot represents one DMR, significantly correlated with age DMRs are shown in red.

To understand global age-associated changes in methylation profiles, we first compared average levels of cytosine methylation within donors of the two age groups. While no difference was observed in overall methylation of CpGs outside of the CpG islands, methylation within the CpG islands significantly increased in older donors (Figure 3B). PCA and hierarchical clustering of CpG island methylation indicated that samples from the two cohorts clustered separately, confirming the presence of a strong age-associated methylation signature (Figures 3C and 3D). This result is concordant with a previous data showing that CpG islands tend to gain methylation with age^12,42,43^. Another global change that we confirmed was increased variability in DNA methylation levels with age^5,14,44,45^. Indeed, when comparing standard deviations of the CpG island methylation levels between monocytes from the two groups, we observed an age-associated increase in variability (Figure 3E). Finally, we have obtained highly consistent results for the two age groups using both Horvath^38^ and Hannum^37^ models, even though a number cytosines had to be imputed in order to use these approaches (**Figure S5B**).

eRRBS, as opposed to widespread DNA methylation array technology, allows for identification of differentially methylated regions composed of multiple concordantly changing cytosines, which is more likely to identify biologically relevant regions. We used the MethPipe pipeline^46^ to perform a genome-wide comparison of the methylomes between the two groups to identify age-associated DNA-methylation signatures, which yielded 1,160 differentially methylated regions (DMRs) (Figure 3F; **Table S12**). Approximately half of the regions were hypermethylated with age and were significantly enriched in CpG islands (Figures 3F and **S5C**), consistent with our observations on the global level (Figure 3B). Other regions lost methylation with age and were typically located outside the CpG islands. These findings indicated the presence of multiple demethylation events that accompanied healthy human aging and were equally as characteristic as a gain of methylation in CpG islands.

Next, we sought to establish robustness of the identified region-based signature by analyzing the behavior of these regions in publicly available datasets. Two whole genome bisulfate sequencing (WGBS) datasets focused on aging of immune cells were published previously^42,47,48^. In both cases group sizes were limited to 1 or 2 samples per group which precluded statistically-sound discovery of aging signature based on these data. However, the datasets could still be used for validation of aging DMRs identified from our data. First, we compared detected DMRs from our dataset to the dataset for purified classical monocytes from cord blood (n=2) and from venous blood from old donors (n=4, age 60-70 y.o.) available from the Blueprint consortium^47,48^. We find that methylation changes in the DMRs that we identified were highly consistent with differences in the same regions observed in Blueprint dataset. For instance, a DMR located within the GRM2 promoter encompassed 50 CpGs and showed significant difference in average methylation between cohorts based on our eRRBS data (Figure 3G, see red and blue lines showing methylation levels in young and old samples respectively). The bar plots in the bottom panel of Figure 3G shows WGBS data from Blueprint Consortium, highlighting distinct age-associated gain in methylation within the same genomic location. High reproducibility was also a case when all DMRs were considered: we found that the changes in DMRs were consistent (Figure 3G) and highly correlated (**Figures S5D**, Spearman’s correlation coefficient = 0.7, p-value < 2.2e-16) between our and Blueprint datasets. Similar results were obtained when we compared our signature against previously published WGBS samples from a newborn and a 103-year-old centenarian^42^ (**Figure S5E**).

Lastly, we compared our signature to published MESA dataset^5,30^ generated using DNA methylation arrays (**Figure S5F**). Out of 1,160 DMRs, only a minute fraction had 3 or more CpG covered in a widely used Infinium 450k methylation array (used in MESA study), meaning that the signature that we identified could not have been found using array-based profiling techniques (Figure 3H). Still, even despite technological differences, PCAs on cytosines located within the DMRs showed a very clear separation by age in MESA data (Figure 3I, left panel). Methylation of the most cytosines located inside DMRs significantly correlated with donor age, and the directionality of the age-associated methylation changes in the MESA dataset matched differences observed in our dataset (Figure 3I, right panel). *Thus, here we report a novel set of age-associated differentially methylated regions and validate the robustness of this signature across different studies.*

### Age-related loss and gain of methylation follow distinct chromatin clues

To understand relationship between identified age-associated changes in methylation patterns and other chromatin features, we focused on the next layer of epigenetic regulation and characterized five post-translational modifications of histone 3 (H3) tails (H3K4me3, H3K4me1, H3K27ac, H3K27me3, H3K36me3) for monocytes from young and old groups (Figures 4A and 4B; **Table S13**). To generate data for histone modifications for each donor, we optimized the Ultra-Low-Input ChIP-seq protocol^15^ that required only 100,000 cells for robust profiling of chromatin modifications from flash-frozen samples. While the ULI-ChIP-seq protocol allows for robust peak calling within individual samples, the data obtained by this method are considerably more variable between samples when compared to classical ChIP-seq approaches^15^ (**Figures S6A and S6B**). To address this limitation, we developed a new computational approach, called SPAN, inspired by a semi-supervised method described by Hocking et al.^49^ (Figure 4A, **Methods** and **Supplementary material**). In this approach, the user annotates a handful of locations as peaks, valleys or peak shores. These annotations are then used to train a model that is optimal for a singular sample (**Figure S7**). Importantly, SPAN peak calling models are trained for each sample individually, preserving sample independence which is critical for downstream comparative analysis. We used our novel approach to analyze data for 191 ULI-ChIP-seq experiments that passed initial QC (Figure 4B). Peaks called by SPAN were significantly more consistent than output of classical peak callers, as shown by various quality control metrics in **Figure S5**. We also evaluated overlap between consensus peaks and existing ChromHMM annotation for CD14^+^ monocytes to show that generated chromatin profiles were consistent with established functional roles of the profiled histone marks^50^ (Figure 4C).

**Figure 4:**
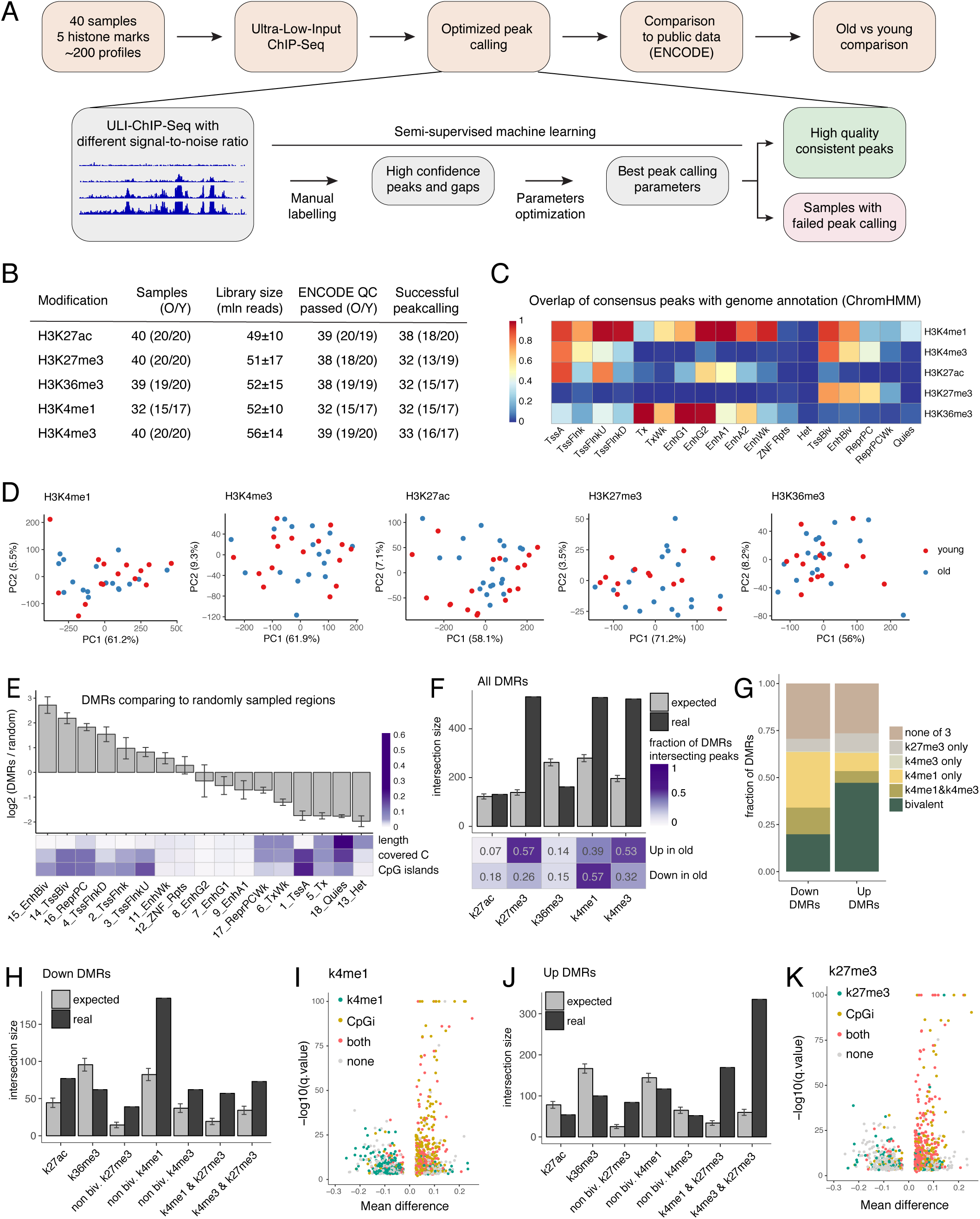
Distinct chromatin signatures of up and down age-associated DMRs. **(A)** Overview of chromatin profiling and processing approach based on semi-supervised machine learning. **(B)** QC characteristics for each chromatin mark. **(C)** SPAN weak consensus peaks (union of peaks confirmed by at least two samples) overlap with ENCODE 18-states ChromHMM markup for CD14^+^ monocytes. Overlap of ChromHMM state with histone mark is shown (overlap of column with row). **(D)** PCA of library depth normalized coverage in weak consensus peaks. Each dot represents one sample, percentage of variance explained by principal components is shown in brackets. **(E)** Enrichment of DMRs in chromatin state segments from ENCODE ChromHMM partition. Upper: bars represent log2 of fold change between real number of DMRs intersecting each chromatin state and expected number of intersections estimated by 100,000 random sampling from genome regions covered in the eRRBS methylation profiling. Error bars show standard deviation (SD). Bottom: heatmap shows general statistics for each chromatin state. Values are normalized within rows. **(F)** Enrichment of all DMRs in genome regions marked by 5 profiled histone modifications. Real number of DMRs intersecting at least one peak from the corresponding mark weak consensus (dark grey bars) was compared to the expected intersection estimated as in 4E (light grey bars). Heatmap below shows fraction of hyper- and hypo-methylated DMRs that co-localize with corresponding histone modification. **(G)** Bar plot represents fractions of the hyper- and hypomethylated DMRs that are marked by different combinations of H3K4me3, H3K4me1 or H3K27me3. Bivalent state refers to regions that are marked by H3K27me3 and either H3K4me3 or H3K4me1. **(H)** Analyses as in 4F for hypomethylated DMRs only and including histone modifications combinations. Non-bivalent H3K4me1 and H3K4me3 refers to regions that lack H3K27me3 mark. In the same manner, non-bivalent H3K27me3 lack both H3K4me1 and H3K4me3. **(I)** Volcano plot as in Figure 3F, colored in accord with DMR intersection with epigenetic feature: both H3K4me1 peak and CpG island (red), only CpG island (yellow), only H3K4me1 peak (green) or neither (grey). **(J)** Same as 4H but for hypermethylated regions. **(K)** Volcano plot as described in 4I. Dot color indicated if corresponding DMR intersects both H3K27me3 peak and CpG island (red), only CpG island (yellow), only H3K27me3 peak (green) or neither (grey).

Having ensured accurate and consistent peak calling, we investigated the age-associated changes in our datasets. First, we performed PCA on weak consensus peaks, using ChIP-seq signals normalized to library depth (RPM). PCA plots for each of the histone marks (Figure 4D) showed an absence of global differences between the two cohorts. Furthermore, we applied a number of differential ChIP-seq tools to the data, which resulted in either no differences or a small number of differences that were not reproducible when analyzed by different tools (see **Methods** and **Supplementary Material**). Visual inspection of the presumably most different peaks also confirmed that these locations were not consistently altered with age (**Figure S8**).

Next, we sought to leverage the multi-omics nature of our dataset and asked if chromatin landscape relates to identified age-associated DMRs. We started by comparing DMRs from our data set (Figure 3) to genomic annotation defined by ENCODE chromHMM for CD14^+^ monocytes (Figures 4E, **S9A and S9B**) and found that DMRs were highly enriched in bivalent states and Polycomb-repressed regions. Conversely, heterochromatin and quiescent states were significantly unlikely to host a DMR (Figure 4E). Enrichment analysis of DMRs against consensus peak sets revealed overrepresentation of DMRs in regions marked with H3K27me3, H3K4me3, and H3K4me1 modifications (Figure 4F), with distinct characteristics for up and down DMRs (Figure 4G). Indeed, hypomethylated DMRs (down DMRs) showed the highest enrichment in non-bivalent H3K4me1-marked regions (Figure 4H). These regions were usually located outside the CpG islands (Figure 4I, left half of volcano plot), and the overall H3K4me1 signal was significantly higher in hypomethylated regions (**Figure S9C**). On the contrary, hypermethylated DMRs (up DMRs) typically co-localized with H3K27me3-marked CpG islands (Figures 4J and 4K, right half of volcano plot). Average normalized signal in DMRs also showed that the general H3K27me3 levels were significantly higher in hypermethylated regions compared to hypomethylated regions (**Figure S9C**, see also **Figures S9C and S9D** for the remaining 3 chromatin marks). *Overall, our data demonstrate that despite the evident age-related re-organization of DNA methylation, the histone modifications H3K4me3, H3K4me1, H3K27ac, H3K27me3, H3K36me3 are stable and do not undergo drastic re-arrangement with age in the basal state. At the same time, age-associated changes in DNA methylation are enriched within specific chromatin features, distinct for up and down DMRs.*

### DNA hypomethylation is directly linked to age-associated transcriptional changes

Significant fractions of both, up and down DMRs (~60% and ~40% respectively) were located in gene promoters (Figure 5A). Therefore, we wanted to evaluate regulatory potential of identified regions and their impact on expression level of the corresponding genes. Strikingly, genes associated with up and down DMRs had a notably distinct expression patterns. Genes with hypomethylated promoters obeyed regular distribution of gene expression, while gene with hypermethylated promoters were either not expressed or expressed at low levels in both young and old individuals (Figure 5B). This lack of expression of genes with hypermethylated regions (up DMRs) in their promoters is consistent with the abundance of repressive H3K27me3 mark in up DMRs (Figure 4K). These observations also suggested that hypomethylated DMRs (down DMRs) were more likely to have a functional impact on gene expression than hypermethylated DMRs (up DMRs). Specifically, we reasoned that age-related hypermethylation of promoters associated with up DMRs could only further decrease expression of corresponding genes that are already lowly expressed, indicating unlikely functional role of this decrease. However, genes associated with promoters that lose their DNA methylation with age (down DMRs) would increase the expression levels of corresponding transcripts, indicating possible functional impact of these regions (Figure 5C, left panel). To test this hypothesis, we determined whether genes with hypo- or hypermethylated DMR intersecting their promoter region ([-10kb; +3kb] around TSS) were significantly enriched within up- or downregulated genes derived from highly-powered MESA transcriptomic dataset (Figure 2C). Strikingly, GSEA analysis indeed showed that hypomethylated DMRs were significantly up-regulated with age in MESA cohort, while no significant enrichment was observed for genes with hypermethylated promoter regions (Figure 5C, right panel).

**Figure 5:**
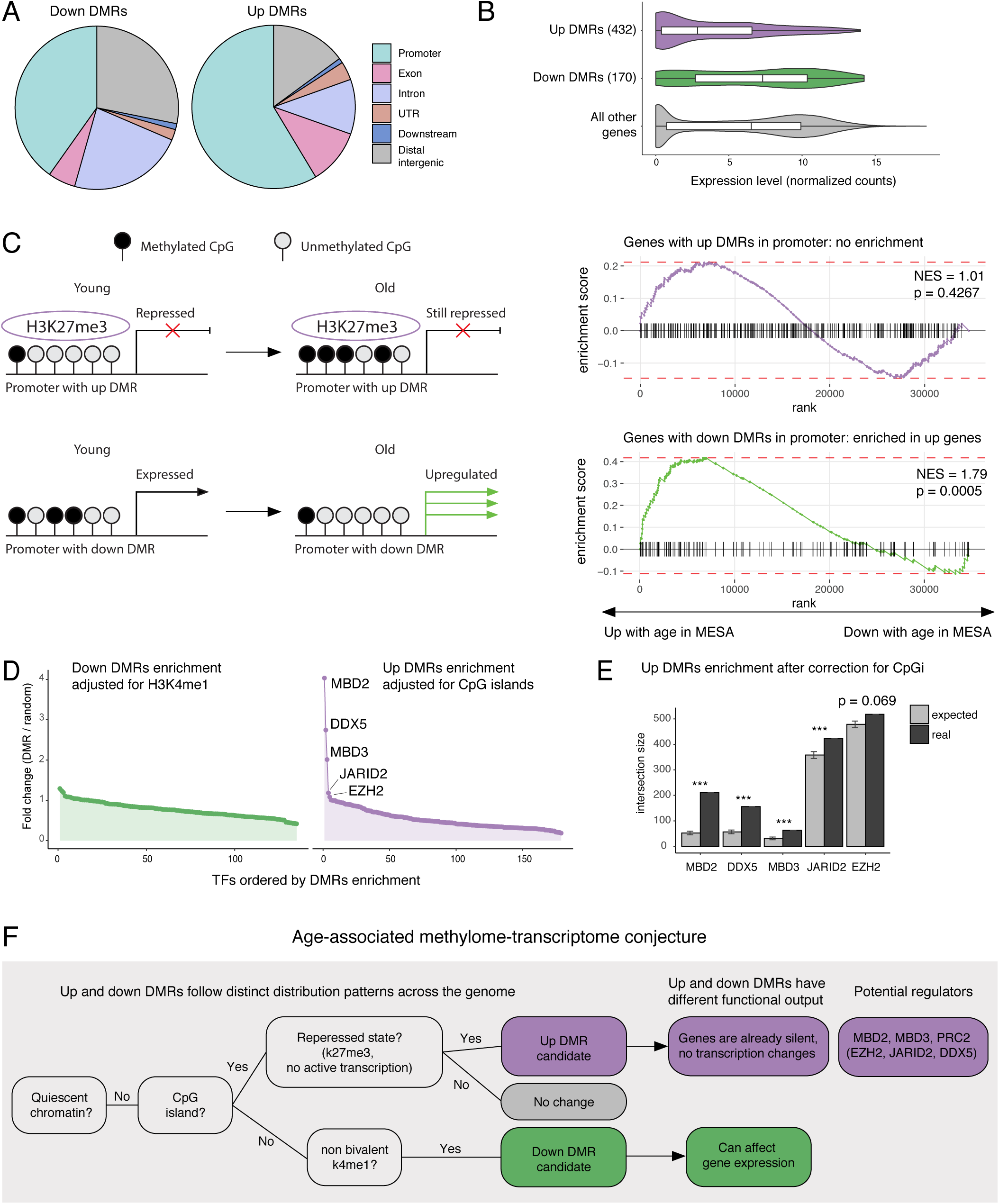
Distinct influence of up and down DMRs on gene expression. **(A)** Pie charts summarize location of hypomethylated (down DMRs, left) and hypermethylated (up DMRs, right) and DMRs in genome with respect to gene annotation. Promoter was defined as [-10kb; +3kb] around TSS. **(B)** Violin plots compare normalized expression levels of genes with hypomethylated regions within promoters, hypermethylated regions within promoters and no methylation changes in promoters. Promoter was defined as in 5A. **(C)** Left: schematic summary of DMR regulatory outcomes hypothesis. Right: GSEA enrichment curves against MESA dataset for genes associated with up and down DMRs as in 5A. **(D)** Enrichment of DMRs in transcription factors (TFs) binding sites annotated by ReMap. Each dot represents a TF. **(E)** Top 5 TF enrichment hits for up DMRs. Barplots compare expected and real intersections between DMRs and TF binding sites as in 4F. **(F)** Proposed model of age-associated alterations in DNA methylation and consequent transcriptional changes.

Next, we wanted to ask if up and down DMRs tended to colocalize with any specific transcription factors. We performed enrichment analysis comparing DMRs to binding sites of 485 transcription factors annotated by the ReMap atlas^51,52^. Consistently with overall tendency of DMRs to localize within regulatory regions (Figure 4E), we identified multiple significant transcription factors for both up and down DMRs. Many of these associations were driven by epigenetic environment of up and down DMRs that were enriched in CpG islands and H3K4me1 marked sites respectively. To that end we adjusted the analysis for the local chromatin structures in order to filter out transcription factors that were inherently associated with CpG islands or H3K4me1 sites (see **Methods** for details). No significant associations remained after the H3K4me1-based adjustment for down DMRs, while four transcription factors were significantly associated with up DMRs after adjustment for CpG islands (Figure 5D; **Table S14**). Among them, JARID2, DDX5 are known to be associated with Polycomb Repressive Complex 2 (PRC2), which is consistent with colocalization of up DMRs with H3K27me3 mark. But surprisingly, we also identified two novel transcription factors from one family – MBD2 and MBD3, that had not been linked to age-related DNA hypermethylation previously (Figure 5E). All detected proteins were highly expressed in monocytes (**Figure S9E**), making them strong candidates for potential regulators of age-associated DNA methylation gain.

Together with enrichment of the DMRs in the specific chromatic signatures, these comparisons highlighted a distinct regulation as well as functional importance of the DMRs identified in our analysis. We propose the following hypothesis to describe the relationship between age-related DNA methylation, its regulation, and its physiological impact on cellular function (Figure 5F)*. Hypermethylated regions are enriched within CpG islands and are often characterized by bivalent signal. These regions are associated with silent genes, and, consequently, their hypermethylation has little impact on the already absent expression. In contrast, hypomethylated regions (down DMRs) are generally outside of CpG islands and enriched in regions marked by H3K4me1 alone. Down DMRs are associated with age-related upregulation of their corresponding genes.*

### Up and down DMRs behave distinctly in different physiological and clinical contexts

Next, we investigated behavior of up and down DMRs in a variety of physiological and clinical settings related to aging. Since most of the public datasets were generated using DNA methylation array technology, we were able to retrieve methylation levels of only a fraction of DMRs in this analysis, as discussed above (Figure 3H). Nevertheless, we found that average methylation level of cytosines from up and down DMRs that were covered by the array was able to accurately capture the difference between young and old twin pairs^53^ (Figure 6A). Furthermore, this metric correlated significantly with age of donors from MESA dataset (Figure 6B, p-value < 0.001 for both), indicating that methylation of these regions continued to change with age even in a population that was older than our studied cohort. However, the behavior of up and down DMRs was different when we looked at the data from bulk brain tissue of a healthy aging cohort^54^: average methylation of up DMRs was still associated with age, down DMRs failed to show significant correlation with age (Figures 6C). While more comprehensive analysis of various cell types and tissues is required to make a solid conclusion, this observation suggests differential tissue/cell type specificity of up and down DMRs.

**Figure 6:**
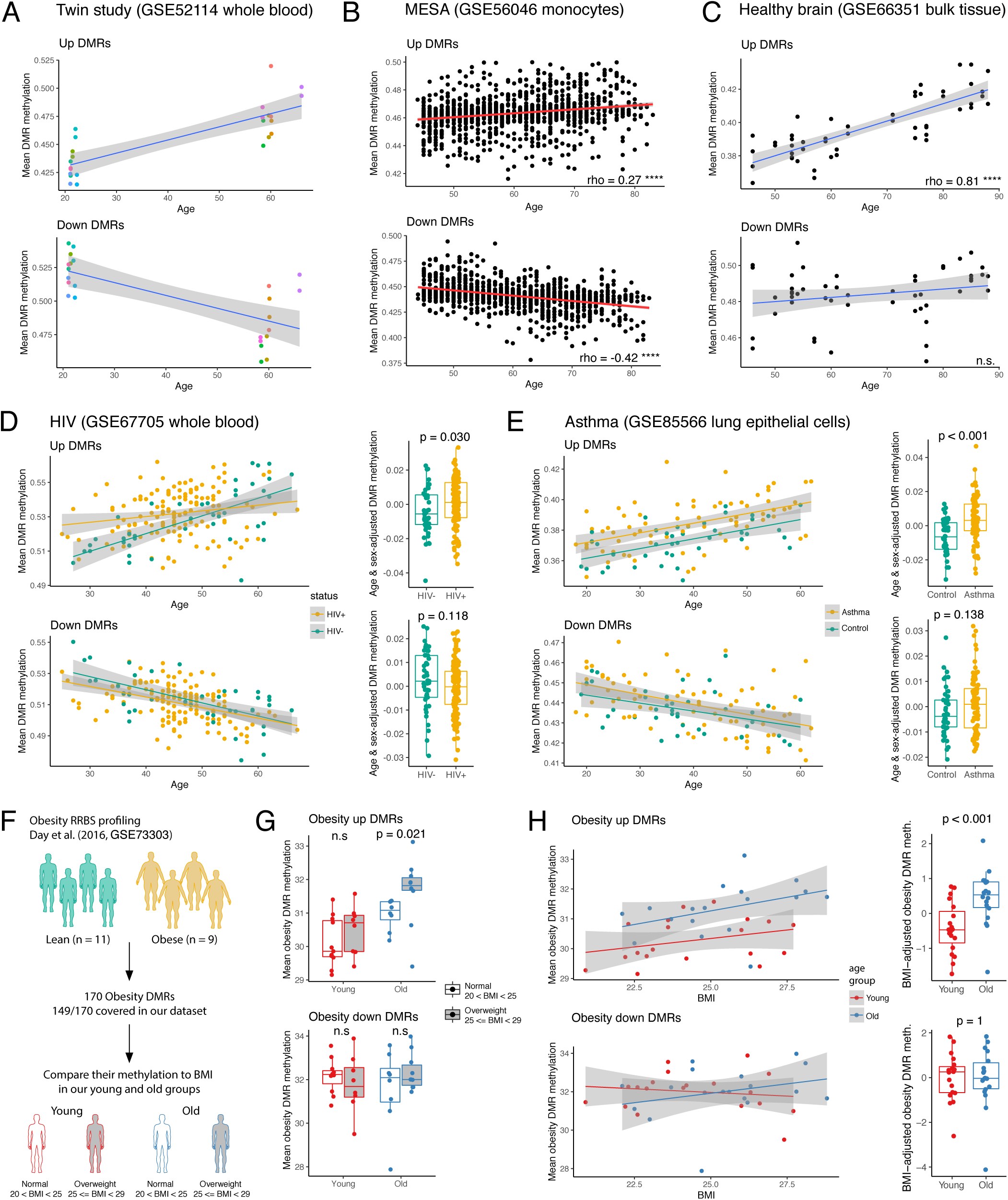
Up and down DMRs behave differently in various pathological contexts. **(A)** Methylation of CpGs that resided in up or down DMR and were profiled by methylation array was averaged for each donor and plotted against donor age. Each dot represents one donor from twin study (color indicates a pair of twins). **(B)** Mean methylation of up and down DMRs as in 6A for samples from **(B)** MESA cohort and **(C)** bulk brain from healthy donors. Spearman correlation coefficient is shown for both. **** indicates p-value < 2.2.e-16, n.s. = nonsignificant correlation. **(D)** Left: plot as 6A for HIV positive (yellow) and negative (green) donors. Right: comparison of age- and sex-adjusted DMR mean methylation between HIV positive and negative donors. Each dot represents one donor. P-values calculated using two-sided Mann-Whitney U test. **(E)** Same as 6D for asthma cases and controls. **(F)** Scheme of analysis of established previously obesity DMRs in our aging cohort. **(G)** Mean methylation of obesity DMRs in our donors split by age and BMI. P-value was calculated using two-sided Mann-Whitney U test. **(H)** Left: mean methylation of obesity DMRs in our donors plotted against donors BMI. Each dot represents one donor. Right: comparison of BMI-adjusted obesity DMR mean methylation between young and old donors. P-values calculated using two-sided Mann-Whitney U test.

To test whether up and down DMRs were relevant for the phenotype of accelerated aging, we compared DMR methylation in HIV positive donors and matching controls^55^. Strikingly, while age was robustly associated with both up and down DMRs, age acceleration in HIV individuals was evident only for hypermethylated up DMRs (Figure 6D, left). Comparison of age- and sex-adjusted levels of mean methylation allowed to quantify this difference, confirming that up DMRs were more methylated in HIV patients irrespective of donors age (Figure 6D, right). Further analysis of published clinical phenotypes revealed novel association with age acceleration: we find that methylome of lungs of asthma patients demonstrate age acceleration in the similar manner as observed for HIV patients (Figures 6E). Up DMRs mean methylation was significantly higher in asthma patients comparing to controls, while down DMR methylation showed no difference. To our knowledge, this is the first report of epigenetic clock acceleration driven by asthma. Notably, we did not observe any age acceleration or delay in either brain or blood of Alzheimer disease patients in these specific regions of DNA, indicating that this effect might be different for either different cell types or DNA regions (**Figure S10A**).

Next, we asked whether changes in DNA methylation driven by lifestyle choices were also affected by age. We focused on obesity and smoking phenotypes due to available RRBS data for these features that enabled investigation of corresponding signature in our dataset. Smoking^56^ did not demonstrate any connection with age-associated DNA hypo- and hypermethylation (**Figure S10C**). However, comparing signatures of obesity reported by Day et al.^57^ to our dataset, we were able to observe a non-trivial relationship between DNA methylation, aging and BMI. Based on their RRBS data Day and co-authors identified 170 obesity DMRs that were well covered in our eRRBS data as well (Figure 6F). Importantly, obesity and age-associated DMRs were distinct sets of genomic regions, and no change in methylation in age-associated DMRs between lean and obese groups was observed (**Figure S10B**). Next, we looked at the obesity-associated DMRs and leveraged variability of BMI in our cohort to compare it to our data. Even though none of our donors were obese, we saw consistent increase in mean methylation of obesity up DMRs in overweight group, but only in case of older donors (Figure 6G). Consistently, when mean methylation of obesity-associated up and down DMRs was plotted against BMI, it was evident that obesity up DMRs were significantly more methylated in the older donors from our cohort (Figure 6H). This suggests that methylome of healthy young individuals is more robust to small variations in lifestyle (e.g. weight change), while in old individuals even mild BMI differences can impact the DNA methylome.

### Hypomethylated regions (down DMRs) are genetically linked to a number of conditions

Lastly, we compared age-associated differentially methylated regions against medical and population genetic databases, such as UK Biobank. We evaluated overrepresentation of DMRs in sets of single nucleotide polymorphisms (SNPs) linked to 34 clinical phenotypes through genome-wide associated studies (GWAS). To evaluate statistical significance and account for incomplete genomic coverage of eRRBS data, we used the simulation procedure defined above for enrichment analysis against histone modifications. While overall distribution of DMRs across the chromosomes was fairly random, we found DMRs to be significantly enriched in four sets of SNPs (Figures 7A and 7B). Only down DMRs showed enrichment in physiological phenotypes (Figure 7B), which was consistent with our previous observation that hypomethylated regions had more functional potential in contrast to hypermethylated DMRs, associated with predominantly silent genes. Specifically, down DMRs were significantly overrepresented in SNPs associated with asthma, level of glycated hemoglobin (Hb), total protein in blood, and multiple sclerosis (MS). Majority of SNPs associated with significant phenotypes were located within human leukocyte antigen (HLA) locus on 6^th^ chromosome (Figures 7C and 7D). This genomic region contains multiple genes regulating immune response, including genes that encode antigen processing and presentation complexes. We confirmed statistically significant enrichment of down but not up DMRs in HLA locus by random simulations (Figure 7E). One of down DMRs resided directly in an exon of HLA-DQB1 gene, encoding a part of MHC class II complex (Figure 7F). Therefore, age-associated changes in DNA methylation have potential to alter antigen presentation by monocytes and monocyte-derived macrophages.

**Figure 7:**
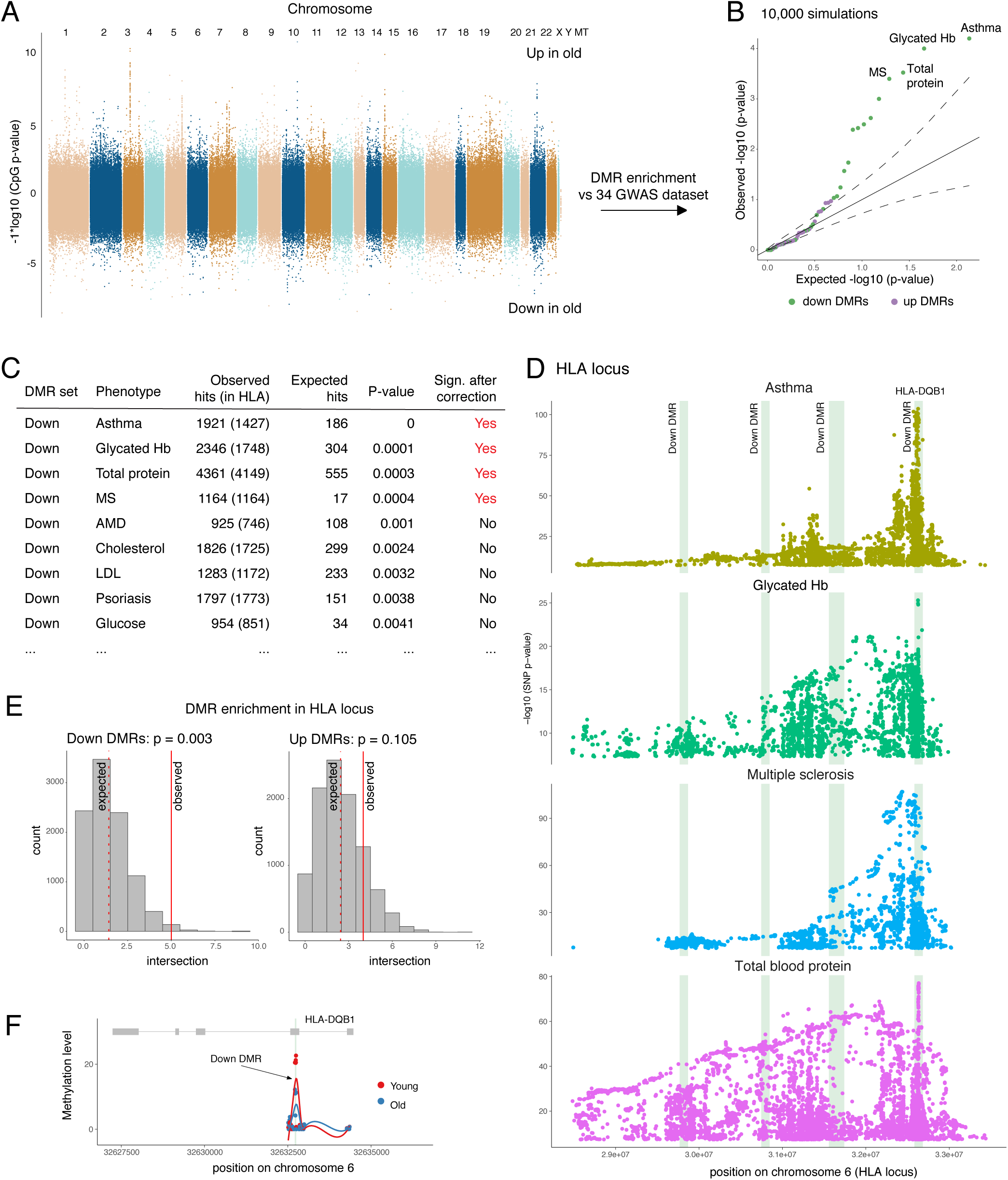
Down DMRs are enriched in genetic susceptibilities loci for asthma, MS, glycated Hb, and total protein. **(A)** Manhattan plot of chromosomal locations and corresponding −log10(p values) for CpGs compared between old and young donors. **(B)** Quantile-quantile plot for DMR enrichment analysis against GWAS datasets. Each dot represents a phenotype compared against down (green) or up (purple) DMRs. **(C)** Top hits of DMR enrichment analysis. **(D)** Manhattan plots of SNPs associated with significant phenotypes and located within HLA locus. Green vertical bars highlight down DMRs located in this locus. **(E)** Enrichment of DMRs in HLA locus. Histogram shows distribution of simulated intersection sizes (10,000 random simulations). Solid red line – real intersection, dashed line – expected intersection. (F) Visualization of DMR intersecting HLA-DQB1 exon. Each dot represents a CpG, green vertical bar highlights the DMR.

## DISCUSSION

We generated and analyzed data describing transcriptomic, proteomic, and epigenetic changes in human CD14^+^CD16^−^ monocytes during physiological aging using a stringently selected healthy cohort. Together with the cell count data, plasma metabolomics and proteomic profiling described in this work, our dataset provides a comprehensive resource obtained in highly controlled settings. We show that in the absence of other inflammatory conditions, aged monocytes are associated with cell intrinsic alterations in DNA methylation patterns, yet are not associated with any dramatic rearrangements in their transcriptional or chromatin profiles for five common chromatin marks (H3K4me3, H3K27me3, H3K4me1, H3K27ac, and H3K36me3). This result is consistent with the absence of a global change in the abundance of various histone modifications in monocytes that was recently shown using CyTOF^14^. Importantly, co-analysis of our data with the MESA transcriptional dataset that includes 1,200 monocyte samples^31^ showed that while transcriptional differences definitely accompanied aging, their magnitude was very small and required high statistical power to be detected, at least in the case of classical circulating monocytes. It is feasible that larger cohort studies might also uncover statistically significant differences in histone modifications between ages, yet it is fair to conclude that the absolute magnitude of such changes would likely be very moderate.

Utilization of next generation sequencing based technology (eRRBS) allowed us to establish a pattern of age-associated changes in DNA methylation and define differentially methylated regions that change with age. In contrast to previous studies that focused on DNA methylation array data and, therefore, identified sets of single-standing CpGs associated with age^5^, we identified age-associated regions that represented concordant changes across multiple neighboring cytosines. This strategy yielded highly robust and physiologically relevant regions as shown by their ability to clearly capture age-specific variations in multiple independent published datasets (Figures 3 and 6). The region-based aging methylation signature identified in this study provided novel insights into age-related DNA methylation events. We found that hypo- and hypermethylation regions were nearly equally frequent in our data and that they differed significantly in their genomic locations, chromatin profiles and relation to the transcriptional activity. Hypermethylated regions were found to be strongly associated with H3K27me3-marked CpG islands residing in promoters of silenced genes. This pattern matches a previously proposed hypothesis that Polycomb is involved in age-associated hypermethylation^58,59^. In addition, we identified MBD2 and MBD3 as new putative regulatory candidates strongly associated with up DMRs. We found hypomethylated DMRs (down DMRs) to be highly enriched in non-bivalent regions carrying the H3K4me1 mark, suggesting their indirect regulatory function and providing a new insight into possible mechanisms of age-associated methylation loss that are yet to be revealed. Unlike CpG islands, the H3K4me1 modification mark sites that often co-localize with cell type specific regulatory regions^60^. Accordingly, this suggests that down DMR might be more cell type specific. Altogether, these observations underscore the existence of orthogonal processes (global and cell-type specific) establishing age-related DNA methylation patterns as well as the importance of profiling pure cell populations.

Most importantly, hypo- and hypermethylated DMRs also exhibited a striking difference in their regulatory impact on transcriptional activity of the associated genes. While genes associated with hypomethylated regions showed a normal expression level distribution, hypermethylated regions were predominantly linked to repressed genes. Accordingly, since methylation gain would only lead to further decrease in transcription, we observed no downstream transcriptional effect of the hypermethylated regions. The hypomethylated regions, however, showed significant enrichment among the genes upregulated with age as defined through re-analysis of data from large independent cohort (MESA). This observation establishes the functional output of age-associated hypomethylation of DNA, proposes that these down DMRs have a more direct effect on the transcriptional state of the cell, and explains the mechanism behind upregulation of a subset of age-related genes. With respect to age-associated hypermethylation of already silenced genes, this process can serve as a protective mechanism that allows cells to avoid tumorigenic transformation^61^.

Enrichment of identified hypomethylated DMRs among functionally important regions further underscores their regulatory potential. We observed overrepresentation of down DMRs in HLA region, suggesting effect of methylation changes on antigen presentation. This enrichment also drives significant intersection of down DMRs with SNPs associated with asthma, multiple sclerosis, and other phenotypes.

Gradual nature of age-associated changes of DNA methylation might explain why diseases such as multiple sclerosis and adult onset asthma develop later in life. Interestingly, in case of asthma, only half of SNP-DMR intersections are explained by demethylation of HLA locus, revealing more robust link between DNA methylation and the disease. Additionally, we showed for the first time that asthma was associated with acceleration of methylation clock in lung epithelial cells.

Overall, we propose a model that separates and characterizes two distinct types of DNA methylation changes and dissects their input into age-associated alterations of cellular state. As next steps, it will be important to understand the functional drivers that control age-associated loss and gain of DNA methylation as well as the higher-level physiological consequences of these changes.

## ACKNOWLEDGEMENTS

The study was supported by funding from the Aging Biology Foundation to Artyomov laboratory and partially by GM125504 to Bagaitkar laboratory and Richard J Lamont. Oltz laboratory is partially supported by ICTS/CTSA Grant# AI134035 and CA188286 from the NCRR. Dixit lab is supported in part by NIH grants P01AG051459, AI105097, AG051459, AR070811, the Glenn Foundation on Aging Research and Cure Alzheimer’s Fund. This publication is solely the responsibility of the authors and does not necessarily represent the official view of NCRR or NIH. We thank the Genome Technology Access Centre in the Department of Genetics at Washington University School of Medicine for help with genomic analysis. The Centre is partially supported by NCI Cancer Centre Support Grant #P30 CA91842 to the Siteman Cancer Centre and by ICTS/CTSA Grant# UL1TR000448 from the National Centre for Research Resources (NCRR), a component of the National Institutes of Health (NIH), and NIH Roadmap for Medical Research. We also wish to thank the Epigenomic Core of Weill Cornell Medicine for the initial analysis of the methylation data (eRRBS and raw data pre-processing). We acknowledge the ENCODE consortium and the ENCODE production laboratories that generated the data sets used in the manuscript. We thank Irina Miralda for figure 1 schematic.

## AUTHOR CONTRIBUTIONS

Conceptualization: I.S., J.B., O.S., M.N.A. Investigation: I.S., J.B, O.S., E.L., S.P., E.W., P.C., G.D., D.A.M., K.Z., S.S., C.C., M.B., L.A., A.S., A.D. E.K., P.T., R.C. V.D.D., M.J., S.D., S.S., E.M.O., M.N.A. Writing-Original Draft: I.S., J.B., O.S., M.N.A. Software: O.S., A.D., E.K., P.T., R.C., S.D.

## METHODS

Detailed protocols and computational methods are available in **Supplementary Material** and on the website http://artyomovlab.wustl.edu/aging/methods.html.

### Ethics Statement

All human studies were approved by the Washington University in St. Louis School of Medicine Institutional Review Board (IRB-201604138). Written informed consent was obtained from all participants. Healthy, Caucasian, non-obese (BMI under 30) males were enrolled in the study in two groups (Figure 1A). Young donors between 24 - 30 years (n=20) and old non-frail donors between 57 - 70 years (n=20) of age were included. Using a screening questionnaire, subjects were asked about lifestyle and health issues. Subjects with any previous history of cancer, inflammatory conditions (rheumatoid arthritis, Crohn’s disease, colitis, dermatitis, fibromyalgia or lupus) or infections (HIV, hepatitis B, C) were excluded. Smokers were also excluded. Blood (~100ml) was collected by venous puncture in the morning (8 – 10 a.m.) after overnight fasting. Some of the older donors self-reported blood pressure alterations/medication, which was not considered as a reason for exclusion.

### Complete blood profiles (CBCs) and monocyte isolation

Briefly, venous blood was collected in Sodium-Heparin vacutainers. CBCs and blood cell differentials were determined using a Hemavet 950FS analyzer within 2 hours of blood draw. Plasma and blood cells were separated using Histopaque-1077 according to the manufacturer’s protocol (SIGMA). Briefly, whole blood was diluted 1:1 with sterile DPBS-2mM EDTA and overlaid (30ml) on to 10ml of Histopaque-1077. Gradients were centrifuged at 500xg for 30 minutes. The upper phase containing plasma was carefully aspirated and stored at −80°C until further use. Peripheral blood mononuclear cells (PBMCs) were isolated and CD16^+^ cells were depleted using anti-human CD16 Milteyni magnetic beads using manufacturer’s protocol. After CD16 depletion, CD14^+^ monocytes were purified using anti-human CD14 magnetic beads (Milteyni). Purity (>98%) was determined by flow cytometry (**Figure S4A**) and cells were either cryopreserved in Cryostor preservation media or snap frozen and stored at −80°C. The detailed protocol is available on the website.

### Flow cytometry

Purified monocytes were incubated with human Fc block and then stained with anti-human CD14 and CD16 antibodies. Data were collected on FACS Canto (BD Biosciences, San Jose, CA) or Cytek-modified FACScan (BD Biosciences and Cytek Development, Fremont, CA) instruments and analyzed with FlowJo (Tree Star, Ashland, OR).

### Bioplex and ELISA

Plasma was profiled using Mesoscale V-plex pro-inflammatory panel I human kit (IFN-γ, IL-1β, IL-2, IL-4, IL-6, IL-8, IL-10, IL-12p70, IL-13, TNF-α) and statistical differences in data obtained were determined by two-sided Mann-Whitney U test. ELISA kits were used to determine GDF-15, Sclerostin (RnD), Osteomodulin (Aviva), Notch1 (Thermofisher), and sCD86 (Abcam) as per manufacturer’s instructions.

### Plasma Profiling: metabolomics

For each donor 250μL of frozen plasma was shipped on dry ice to Metabolon (http://www.metabolon.com) for Liquid Chromatography-Tandem Mass Spectroscopy (LC-MS). 734 peaks were annotated. Pre-processing was performed by Metabolon: peaks were quantified using area-under-the-curve and each compound was corrected in run-day blocks by registering the medians to equal one and normalizing each data point proportionately (termed the “block correction”) to correct variation resulting from instrument inter-day tuning differences. YD20 was excluded from the analysis as an outlier based on PCA. Significant differences in metabolites between cohorts were determined using two-sided Mann-Whitney U test. Pathway annotation was provided by Metabolon. For each pathway collective statistical significance was determined by comparing mean log2FC of pathway members to zero using a one sample two-sided Mann-Whitney U test. P-values were adjusted for multiple testing using the Benjamini & Hochberg method. Significance threshold was set to 0.05 in for all comparisons.

### Plasma Profiling: proteomics

For each donor 500μL of frozen plasma was shipped on dry ice to the GTAC core facility at Washington University in St. Louis for high-density protein expression analysis via SomaScan assay^62^. Profiles for ~1,300 analytes were acquired. Pre-processing was performed by GTAC core facility at Washington University in St. Louis: raw RFU measurements for every SOMAmer reagent were normalized subsequently with hybridization normalization, plate scaling, median scaling, and calibrator normalization and transformed in log2 scale. Differential analysis was done for young vs. old groups using quantile-normalized data for all 40 samples. For differential analysis, functions lmFit and eBayes from the Limma package (version 3.34.5)^63^ were used. P-values were adjusted for multiple testing using the Benjamini & Hochberg method.

Correlation analysis (**Figure S3E**) used significant plasma proteins and all profiled plasma metabolites that had non-zero variance. Pheatmaps of absolute values of Spearman correlation coefficients were plotted using Pheatmap R package.

### RNA-Seq data: young versus old monocytes

Total RNA was isolated from snap frozen monocyte pellets using Qiagen’s RNeasy Mini Kit according to the manufacturer’s protocol. RNA concentration and integrity were assessed by Agilent 2200 Tape Station. RNA samples were submitted to BGI (Hong Kong, China) for long non-coding RNA sequencing and small RNA sequencing. After library construction, paired-end 100 base pair reads were generated on the DNBseq platform.

Fastq files for each sample were aligned to the hg19 genome (Gencode, release 28) using STAR (v2.6.1b) with the following parameters: STAR --genomeDir $GENOME_DIR --readFilesIn $WORK_DIR/$FILE_1 $WORK_DIR/$FILE_2 --runThreadN 8 --readFilesCommand zcat -- outFilterMultimapNmax 15 --outFilterMismatchNmax 6 --outReadsUnmapped Fastx -- outSAMstrandField intronMotif --outSAMtype BAM SortedByCoordinate -- outFileNamePrefix. /$^62^. Quality control for each sample was performed by FastQC (v0.11.3) and Picard tools (v2.18.4). Quantification was done using htseq-count function from HTSeq framework (v0.9.1): `htseq-count -f bam -r pos -s no -t exon $BAM $ANNOTATION > $OUTPUT`.

Raw counts were normalized prior to PCA via getVarianceStabilizedData function from DeSeq2 package (v1.24.0). Differential expression analysis was done using DESeq function from DeSeq2 with default settings. Following design was used in the analysis: *gene ~ age + batch + PC1 + PC2 + PC3*. PC1, PC2, and PC3 are three main principle components explaining genetic variability in the cohort^64^. Genotype data was retrieved from ChIP-seq raw reads as described in **Supplementary Materials** and **Figure S11**. Significance threshold was set to adjusted P-value < 0.05.

### Publicly available transcriptional data

Transcriptomic dataset for MESA cohort was re-analysed^31^. GSE56045 contains normalized expression values for purified human monocytes. Differential analysis was performed using Limma package (v3.26.5). We accounted for cofounding variables by including chip and race-gender-site parameters into model design: *gene ~ age + chip + race-gender-site*. Age was used as continuous variable, no separation into age groups was performed. P-values were adjusted for multiple testing using Benjamini & Hochberg method. Significance threshold was set to FDR < 0.05. Gene set enrichment analysis via fgsea R package^65^ (v1.10.0) was used to identify enriched pathways and plot enrichment curves.

### RNA-Seq data: differentiation and stimulation of monocytes

Differentiation of primary human CD14^+^CD16^−^ monocytes into resting macrophages occurred after 7 days of culture in RPMI media supplemented with 11 mM glucose, 2 mM glutamine and 10% fetal calf serum in the presence of 50 ng/ml of M-CSF (Peprotech, cat# 300-25). 5 × 10^5^ CD14^+^CD16^−^ monocytes or × 10^5^ resting macrophages were activated with 10 ng/ml of lipopolysaccharides (LPS) from E. coli (Sigma, O111:B4) or mock for 24 hr.

RNA was isolated from monocytes and macrophages using AllPrep DNA/RNA Mini Kit (Qiagen, cat# 80204) and treated with RNAse-free DNAse (Qiagen, cat# 79254). Libraries were prepared as described previously^66^. Briefly, cDNA was synthesized using custom oligo(dT) primers with a barcoded adaptor-linker sequence (CCTACACGACGCTCTTCCGATCT-XXXXXXXX-T15). Barcoded cDNA was pooled together based on ActB qPCR values and the RNA-DNA hybrids were degraded by consecutive acid-alkali treatment. A second sequencing linker (AGATCGGAAGAGCACACGTCTG) was ligated via T4 ligase (NEB), followed by clean up with SPRI beads (Beckman-Coulter). The libraries were amplified by 12 cycles of PCR and cleaned up with SPRI beads, yielding strand-specific RNA-seq libraries. Data were sequenced via HiSeq 2500 40 bp x 10 bp paired-end sequencing.

Files obtained from the sequencing center were demultiplexed using fastq-multx tool. Fastq files for each sample were aligned to the hg19 genome (Gencode, release 28) using STAR (v2.6.1b) with the following parameters: STAR --genomeDir $GENOME_DIR --readFilesIn $WORK_DIR/$FILE -- runThreadN 8 --outFilterMultimapNmax 15 --outFilterMismatchNmax 6 -- outReadsUnmapped Fastx --outSAMstrandField intronMotif --outSAMtype BAM SortedByCoordinate --outFileNamePrefix. /$^62^. Quality control for each sample was performed by FastQC (v0.11.3) and Picard tools (v2.18.4). Aligned reads were quantified using a HTSeq-based quant3p script^67^ (available at https://github.com/ctlab/quant3p) to account for specifics of 3′ sequencing: higher dependency on good 3′ annotation and lower level of sequence specificity close to 3′ end. DESeq2 (v1.24.0) was used for analysis of differential gene expression.

### Monocyte proteomic data

For monocyte proteomic analysis we used an independent cohort of donors that was recruited using the same inclusion criteria as in the rest of the study (n=11 samples per group). For each donor, snap frozen monocytes (1×10^6) were shipped on dry ice to Biognosys for protein extraction and LC-MS analysis. Details of Biognosys Discovery protein profiling pipeline (HRM ID+ mass spectrometry) can be found on their website (https://biognosys.com/technology/#discovery-proteomics). On average, 5,580 proteins were quantified per sample. In total, 5,804 proteins represented by 74,348 peptides were quantified across all samples. Statistical assessment was performed by Biognosys (**Table S9D**). In brief, for each protein the fold change of each peptide ion variant was estimated as average abundance of peptide ion variant across biological replicates in old group / average abundance of peptide ion variant across biological replicates in young group. The values then were log-transformed and fold changes of all peptides belonging to the same proteins were compared to zero using two-sided paired t-test. Multiple testing correction was performed as described in Storey et al.^68^ We removed MHC proteins from the analysis due to high similarity between the variants that could not be accurately resolved using this data. For visualization purposes (Figure 2G) we used protein intensities estimated as an average of the top three peptides for each protein (**Table S9E**).

### Monocyte eRRBS data acquisition

Genomic DNA was extracted from snap frozen monocyte pellets (1-2 million cells) using the Qiagen’s all prep RNA/DNA extraction kit as per manufacturer’s instructions. Fragment size was confirmed by Agilent Tape Station to be >40 kb with no significant degradation. 1μg of genomic DNA was submitted to the Epigenomics Core at Weill Cornell Medicine for eRRBS library prep and sequencing.

### Monocyte eRRBS data analysis: initial processing, QC and filtration

Initial processing of raw data was performed by Cornell Epigenomics Core according to the standard pipeline described in^41^. Average conversion rate was higher than 99.75% for all samples and sequencing depth varied between 80,359,782 and 58,727,376 reads per sample (**Figure S5A, Table S11**). The eRRBS protocol mainly focuses on CpG-rich regions so only a fraction of the entire genome was covered. Therefore, while cytosines in CpG, CHH and CHG contexts were present in the dataset, only cytosines in the context of CpG were well-covered across the majority of samples. Thus, we focused on DNA methylation in CpGs only (**Figure S5G**). Overall, 20,077,756 CpGs were covered in at least one sample. More than 500,000 cytosines were covered with at least 10 reads in each of samples, in accord with field standards (Fig. 3A)^41^. We removed all cytosines with insufficient coverage: average coverage across all 40 samples was required to be greater than or equal to 10 reads. 2,808,448 CpG cytosines remained in the analysis after filtration (we refer to this set as “covered CpG cytosines”, and all the downstream analysis used only these cytosines). Overall, for approximately 84% CpG islands at least one cytosine in CpG context was covered in the experiment. CpG island annotation was downloaded using UCSC Table Browser for hg19^69^.

Exploratory data analysis was performed to verify data quality and remove outliers that had skewed methylation patterns compared to the majority of the donors. PCA revealed 3 samples that were distinct from the entire set of donors (**Figure S5H**). This was further confirmed by analysis of methylation distribution and hierarchical clustering (**Figures S5I and S5J**). These samples (1 young donor and 2 old donors) were removed from further analysis. Methylation data was acquired in two separate batches with similar library preparation protocols (10 young and 10 old in each batch). Therefore, we account for a possible batch effect (**Figure S5K**) in all the following analysis.

To apply Hannum^37^ and Horvath^38^ methylation clock models, methylation levels of non-covered in our dataset CpGs were imputed as an average methylation within [-100kb; +100kb] window around a CpG. Methylation was set to zero if imputation was not possible (no CpGs were covered in the window.)

### Monocyte eRRBS data analysis: CpG islands methylation analysis

To investigate global changes in the methylome, we averaged methylation levels of all cytosines in CpG context inside and outside CpG islands for each donor (Figure 3B). 2-way ANOVA *(~ age + batch*) was used to calculate p-values. The mean methylation level of each CpG island refers to a mean of methylation levels of all covered cytosines within the island. PCA on the centered and scaled values and hierarchical clustering via Ward algorithm with Manhattan distances were performed for visualization purposes (Figures 3C and 3D). To compare CpG island variability between two age groups we calculated the standard deviation of mean methylation level for each CpG island within young and old cohorts (Figures 3E). Two-sided Wilcoxon paired rank-sum test was used to calculate p-values.

### Monocyte eRRBS data analysis: differential methylation analysis

We used the Methpipe pipeline (v3.4.3) to find differentially methylated regions (DMRs)^46^. Initial per-donor methylation files were converted to Methpipe methcount format and merged into a proportion table using merge-methcounts function. Design table included intercept, age and batch. For each cytosine linear model was fit by radmeth regression function (`radmeth regression -factor age -o OUTPUT_CYTO DESIGN_TBL PROP_TBL`), significance combining and adjustment for multiple testing was performed by radmeth adjust command (`radmeth adjust -bins 1:50:1 OUTPUT_CYTO > ADJ_CYTO`). Significantly different cytosines were merged into regions by radmeth merge (`radmeth merge -p 0.05 ADJ_CYTO > DMRS`). Resulting regions were subjected to further filtration: number of cytosines within the region ≥ 3 and absolute value of difference ≥ 0.025.

For each DMR, a combined p-value was calculated by Fisher method (function fisher.method from BisRNA R package^70^) from Methpipe p-values for cytosines located within the region. Combined p-values were used for visualization purposes only (volcano plots, Figures 3F, 4I and 4K). Resulting DMRs were annotated using ChIPseeker package^71^. A promoter was defined as [-10kb; +3kb] relative to TSS. To find DMRs that intersect CpG islands we used bedtools2 (v2.25.0) intersect function^72,73^.

### Monocyte eRRBS data analysis: comparison to public datasets

Publically available Blueprint dataset EGAD00001002523 was used for DMRs validation^47,48^. Methylation signal in BigWig format is downloaded from publicly available Blueprint online portal for classical CD14^+^CD16^−^ monocytes from cord blood (n=2) and old donors (n=4, 60-70 y. o.). Bedtools2 (v2.25.0) map function was used to calculate mean methylation level for each DMR in Blueprint samples (bedtools map -a DMRS.bed -b BP_SAMPLE.bedGraph -c 4 -o mean)^72,73^. IGV (v2.3.72) was used for BigWig-file visualization (Figure 3G)^74,75^.

WGBS data for cord blood and a 103 y.o. centenarian previously published by Heyn et al.^42^ was downloaded from GEO database (GEO31263) in BED format. DMR methylation (**Figure S5E**) was estimated as mean methylation of all CpGs residing inside DMR and covered in WGBS sample.

Methylation profile for MESA cohort was downloaded from GEO database (GSE56046). M-values were transformed into beta-values by m2beta function from lumi package^76^ (v2.26.4). Batch effect stemming from chip and race-gender-site variables was removed by ComBat function from sva package (v3.22.0). For Figures 3I and **S5F** we used only methylation levels of CpGs residing within DMRs. In Figure 3I (right) methylation level of each DMR in MESA was estimated as mean methylation of all CpGs profiled by the array that reside within corresponding DMRs (for most DMRs data for only one CpG was available). Spearman correlation between DMR methylation and donor age was calculated.

### Ultra-Low input ChIP-Seq (ULI-ChIP)

Aliquots of 100,000 CD14^+^CD16^−^ monocytes were thawed on ice for 5 minutes then immediately resuspended in 20μL of EZ Nuclei Isolation Buffer (Sigma-Aldrich), and incubated for 5 minutes on ice. Samples were digested using 2 Units/μL MNase in 20μL MNase Digestion Buffer (NEB) for 5 minutes at 37°C. Reactions were stopped by the addition of 10mM EDTA and 0.1% Trition / 0.1% Deoxycholate (final concentration). Isolated chromatin was incubated on ice for 15 minutes, followed by vortexing on a medium setting for 30 seconds. The volume was adjusted to 200μL with Complete IP Buffer (20mM Tris-HCL pH 8.0, 2mM EDTA, 150mM NaCl, 0.1% Triton X-100, 10mM Sodium Butyrate, 1x Protease Inhibitor Cocktail, 1mM PMSF), and incubated for 1 hour at 4°C on a gentle rocking platform. 10% of the total chromatin was removed to assess digestion efficiency and to use as an input control.

Chromatin for immunoprecipitation was pre-cleared using Protein A Dynabeads (Invitrogen) for 1 – 4 hours at 4°C and subjected to IP overnight at 4°C (0.3 µg H3K27me3 Millipore 07-449; 0.05 µg H3K27Ac Abcam ab4729; 0.03 µg H3K4me3 Abcam ab8580; 0.2 µg H3K4me1 Abcam ab8895; 0.1 µg H3K36me3 Abcam ab9050). Bead-chromatin complexes were washed using low-salt wash buffer (0.1% SDS, 1% Trition X-100, 2mM EDTA, 20mM Tris pH 8.0, 150mM NaCl, 1x Protease Inhibitor Cocktail, 10mM Sodium Butyrate), high-salt wash buffer (0.1% SDS, 1% Trition X-100, 2mM EDTA, 20mM Tris pH 8.0, 500mM NaCl, 1x Protease Inhibitor Cocktail, 10mM Sodium Butyrate). Chromatin was eluted from the beads using elution buffer (1% SDS, 100mM NaHCO3) by shaking for 1 hour at 65°C. DNA was then purified by phenol-chloroform extraction using Maxtract tubes (Qiagen), and ethanol precipitated overnight. Immunoprecipitated DNA was prepared for sequencing on the Illumina platform using the NEBNext ChIP-Seq Library Prep Master Mix Set using modified Illumina TruSeq adapters.

### ULI-CHIP-Seq peak calling

ULI-ChIP-Seq data pre-processing consisted of the following steps: (1) QC of raw reads including reads quality, length, duplication rate, GC-content; (2) alignment of the raw reads to human genome build hg19; (3) visual inspection of tracks.

Reads quality control (step 1) showed high quality of the data: read length was 51bp, average duplication level was less than 20%, GC content was about 47%, and average library size was ~50 million reads for all the modifications (Figure 4B). Full QC of the raw data is available in the **Table S13**. Distinct nucleotide sequence was overrepresented in the first 5 bp of the reads in all ULI-ChIP-seq libraries, which was an artifact of the ULI-ChIP-Seq protocol. Therefore, first 5 bp were clipped during alignment step. Reads were aligned on the hg19 reference genome using bowtie (v1.1.1), only uniquely mapped reads were used in the downstream analysis. For details see **Supplementary Material**. ULI-ChIP-Seq data processing pipeline is available on GitHub: https://github.com/JetBrains-Research/washu

While almost all the libraries passed QC, signal to noise ratio varied considerably within the cohort (**Figure S6A**), which is a known issue of the ULI-ChIP-seq data. This variation made application of the “golden standard” peak caller not feasible in case of our dataset (**Figure S6B**). To overcome this problem, we developed a novel semi-supervised peak calling algorithm SPAN, based on the idea proposed by Hocking et al^49^. SPAN preprocesses each sample separately to train underlying statistical model. Next, peak calling parameters are optimized individually for each sample based on a single manually-created markup. Detailed description of SPAN is available in **Supplementary Material**. SPAN can be applied to both ultra-low-input and conventional ChIP-seq datasets, which was shown using several publicly available datasets (see **Supplementary Material**).

We used SPAN to analyze data for 191 ULI-ChIP-seq experiments that passed initial QC (Figure 3B). The number of peaks called by SPAN was significantly more consistent for different donors compared to traditional peak calling approaches (**Figure S6D**), which was further supported by improved overlap between peaks for each pair of donors (**Figures S6C and S6E)**. Each dot in **Figure S6E** indicates a fraction of overlapping peaks between two samples, with ~400 such dots making up the value for each bar.

This representation allows direct comparison of peak calling consistency between available methods and illustrates improvements achieved by SPAN (**Figures S6F and S6G**), which can also be seen from a directional overlap between all the samples, histone modifications, and donors (**Figure S6H**). To ensure that SPAN does not introduce unforeseen artifacts, the resulting peaks were compared to the peaks identified for CD14^+^ monocytes in the ENCODE project using conventional ChIP-sequencing^77^ (**Figure S6I**). In comparison with data from ENCODE we used only male samples (GSM1102782, GSM1102785, GSM1102788, GSM1102793, GSM1102797). For consistency, we took raw data and applied our processing pipeline using the same labels that were generated for ULI-ChIP-Seq data (validity of the markup was confirmed by visual exploration).

### Differential ChIP-Seq analysis

To describe age-associated changes in histone code, we performed a differential analysis of the ChIP-seq data between young and old cohorts. First, we performed PCA on signal normalized to libraries depth (RPM) in weak consensus peaks. Second, we ran differential analysis tools, described in ^78^: DiffBind, MACS2 bdgdiff and diffReps. We also added ChIPDiff to the comparison. Since Macs2 bdgdiff does not support replicated data we pooled samples into a single BAM file. Single pooled control for all young and old samples was used with all tools. See **Supplementary Material** for exact setting used for differential analysis.

### Integration methods: overrepresentation analysis

We used the ENCODE 18 states chromHMM annotation (ENCSR907LCD, CD14^+^ monocytes) for overrepresentation analysis. Since our data was generated using eRRBS technology, only a fraction of the genome was covered, and we had to account for incomplete background in our analysis. First, a BED file with all covered regions was generated by merging filtered CpGs located closer than read length (50bp) and expanding regions to be not shorter than read length. Next, bedtools2 (v2.25.0) function shuffle was used to simulate 100,000 sets of non-overlapping regions with length distribution that matched DMRs one and located within the covered fraction of the genome. For each state we calculated the real number of DMRs intersecting the state as well as intersection size with all simulated sets (`bedtools intersect - a RegionSet -b chromHMMState -u`). The expected size of intersection for each state was estimated as the mean of simulated intersection sizes. Corresponding fold change (DMRs / random) was defined as (real intersection / expected intersection). Significance of over- and under-representation for each chromHMM state was calculated as (number of simulated sets with intersection higher than real / 100,000) and (number of simulated sets with intersection lower than real / 100,000), respectively (**Figure S5D**). Benjamini & Hochberg method was used to adjust for multiple testing.

Overrepresentation analysis against peaksets was performed similarly. Weak consensuses (peaks confirmed by at least 2 samples) for each histone modification profiled in our study were used in this analysis.

PCA of chromHMM states was performed similarly to PCA of CpG islands mean methylation (described in eRRBS methods section): mean methylation level of each region refers to a mean of methylation levels of all covered cytosines within the region. Values were centered and scaled for PCA (**Figure S9B**).

### Integration methods: comparison of ChIP-seq signal

ChIP-seq signal intensities in DMRs were estimated using DiffBind package (v2.4.8). We used counts = dba.count(dba(sampleSheet = sheet), fragmentSize = 125, bRemoveDuplicates = TRUE), counts = dbs.count(counts, peaks=NULL, score=DBA_SCORE_TMM_MINUS_FULL), tmm_normalized = dba.peakset(k27ac_counts, bRetrieve = TRUE, DataType = DBA_DATA_FRAME) commands to retrieve TMM-normalized signal values for DMRs. A two-sided Mann-Whitney U test was performed to calculate p-value between hypo- and hypermethylated DMRs.

### Integration methods: DMR-associated expression patters and enrichment in MESA transcriptomic data

To characterize expression of genes associated with changing methylation we selected all genes with DMRs located in their promoters ([-10kb; +3kb] around TSS, see description of the annotation procedure above). Raw counts were generated for our RNA-seq dataset by HTSeq as described above. Gene expression levels for all gene were log-transformed (log2(1 + raw count)) and quantile normalized.

Same sets of genes were used to perform GSEA using ranked list obtained for MESA transcriptional dataset (see details above). Genes were ordered based on t-statistics generated by limma. 10,000 permutations were performed to evaluate significance via fgsea package^65^ (v1.10.0).

### Integration methods: TF binding sites overrepresentation analysis

Binding sites of 485 transcription factors were obtained from ReMap 2018 (v1.2) website (http://pedagogix-tagc.univ-mrs.fr/remap/index.php?page=download). Hg19 merged peak files were downloaded. General workflow of the overrepresentation analysis is described above. To account for enrichment of up DMRs in CpG islands we used only covered regions that intersect a CpG island to simulate 10,000 random up DMRs. Similarly, for down DMR analysis covered regions that intersect H3K4me1 weak consensus peak were used as background to generate 10,000 random down DMRs.

### Integration with public data (Figure 6)

First, we investigated behavior of up and down DMRs in several DNA methylation array-based datasets. Mean methylation level of up or down DMRs was estimated for each sample in a public dataset as an average of all CpGs covered in the dataset that reside within the corresponding set of DMRs. Spearman correlation coefficients between DMR mean methylation and donors age were calculated when applicable. We fit *methylation ~ age + sex* model using lm and took residuals to estimate age and sex-adjusted DMR methylation. Two-sided Mann-Whitney U test was used to calculate p-values.

We explored seven public datasets generated using DNA methylation arrays:

1. Twin study dataset^53^ (GSE52114). Probe methylation values were used as provided in GEO.
2. MESA dataset^5,30^ (GSE56046). See details of data processing above.
3. Bulk brain tissue dataset (GSE66351). Original study^54^ was focused on Alzheimer’s disease, so we filtered out all affected individuals from the dataset and kept only a cohort of healthy controls. We used data from bulk brain tissue and removed one sample (GSM2808945) as an outlier based on PCA.
4. HIV dataset^55^ (GSE67705). We used whole blood data in our analysis. Following sample were excluded as outliers based on PCA: METI-24, METI-1, METI-8, CHAR23, CHAR123, CHAR86.
5. Asthma dataset^79^ (GSE85577).
6. Alzheimer’s disease glia and neurons dataset^54^ (GSE66351). Two samples younger than 20 were removed from the analysis as outliers (all other donors were over 60 y.o. in both cohorts).
7. Alzheimer’s disease whole blood dataset^80^ (GSE59685).

Second, we compared our data to RRBS-based datasets that focused on obesity and smoking.

1. 170 obesity DMRs were acquired from supplementary table 3 of the original study^57^. 149 obesity DMRs had at least one CpG covered in our data. Mean methylation of up and down obesity DMRs was calculated for all our donors as an average methylation of all CpGs from obesity up/down DMRs that were covered in our data (Figures 6G and 6H). BMI-adjusted methylation of obesity DMRs was calculated by fitting *methylation ~ BMI* model and taking residuals. P-values were calculated using two-sided Mann-Whitney U test.
2. Obesity data was downloaded from GEO (GSE73303). Mean methylation of aging DMRs was estimated as described for array datasets (**Figure S10B**). Three samples were excluded as outliers based on PCA: GSM1890526_T10_Obese, GSM1890518_T06_Lean, GSM1890521_T16_Lean.
3. Smoking DMRs were defined by Wan et al.^56^ (supplementary table 4).

### Integration with GWAS data (Figure 7)

GWAS summary statistics of 34 phenotypes produced by large consortia studies were obtained (Neale lab analysis of UK Biobank, http://www.nealelab.is/blog/2017/7/19/rapid-gwas-of-thousands-of-phenotypes-for-337000-samples-in-the-uk-biobank), only variants with association p-values below 5e-8 were included. Overrepresentation analysis based on 10,000 random up and down DMRs is described above. Here DMRs were additionally flanked by 50kb on each side. With total 68 comparison (34 for up DMRs + 34 for down DMRs), significance threshold was set to 0.05 / 68 = 7.4e-4.

### Data visualization: boxplots

In all figures the lower and upper hinges of all boxplots represent the 25th and 75th percentiles. Whiskers extend to the values that are no further than 1.5*IQR from either upper or lower hinge. IQR stands for inter-quartile range, which is the difference between the 75th and 25th percentiles.

### Data availability

Raw sequencing data are deposited to Synapse repository (deposition number acquisition in process). Processed sequencing data, as well as raw and processed metabolic and proteomics data are available at the dedicated online portal at http://artyomovlab.wustl.edu/aging/.

