## Supplementary figures for "Epigenetic aging of classical monocytes from healthy individuals"

### Figure S1

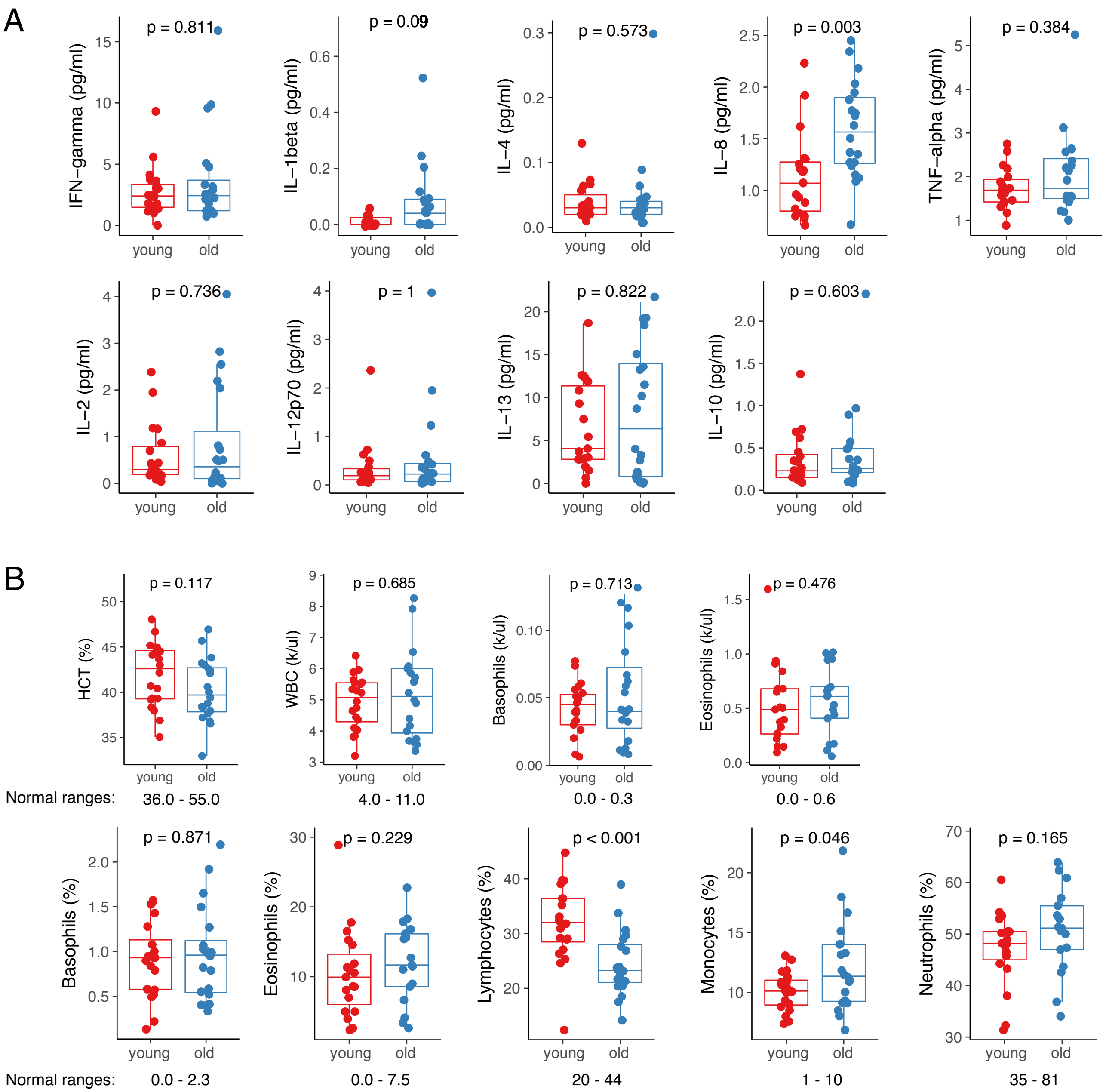
A
p = 0.811
IFN−gamma (pg/ml)
p = 0.09
IL−1beta (pg/ml)
p = 0.573
IL−4 (pg/ml)
p = 0.003
IL−8 (pg/ml)
p = 0.384
TNF−alpha (pg/ml)
p = 0.736
IL−2 (pg/ml)
p = 1
IL−12p70 (pg/ml)
p = 0.822
IL−13 (pg/ml)
p = 0.603
IL−10 (pg/ml)
young
old
B
p = 0.117
HCT (%)
p = 0.685
WBC (k/ul)
p = 0.713
Basophils (k/ul)
p = 0.476
Eosinophils (k/ul)
Normal ranges: 36.0 - 55.0
4.0 - 11.0
0.0 - 0.3
0.0 - 0.6
p = 0.871
Basophils (%)
p = 0.229
Eosinophils (%)
p < 0.001
Lymphocytes (%)
p = 0.046
Monocytes (%)
p = 0.165
Neutrophils (%)
Normal ranges: 0.0 - 2.3
0.0 - 7.5
20 - 44
1 - 10
35 - 81

### Figure S2

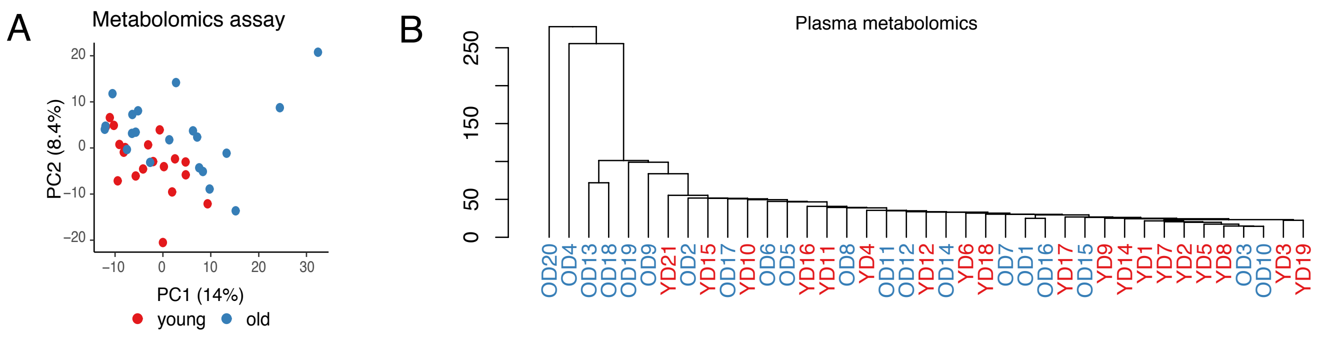


C

Significant metabolities:

21 decrease in old

- 15 steroids
- 1,5-anhydroglucitol
- 5-bromotryptophan
- pro-hydroxy-pro
- glutarate
- 1,2-dilinoleoyl-GPC (18:2/18:2)
- 3beta-hydroxy-5-cholestenoate

18 increase in old

- cystine
- arabitol/xylitol
- N1-methylinosine
- ceramide
- alpha-tocopherol
- 1-(1-enyl-oleoyl)-GPE (P-18:1)*
- N-trimethyl 5-aminovalerate
- 4-acetamidobutanoate
- 3-methylglutaconate
- glucuronate
- formiminoglutamate
- phenylacetylglutamine
- pyridoxate
- 3-hydroxy-3-methylglutarate
- N1-Methyl-2-pyridone-5-carboxamide
- N1-Methyl-4-pyridone-3-carboxamide
- N-acetylglucosamine/N-acetylgalac-tosamine
- 1,3,7-trimethylurate

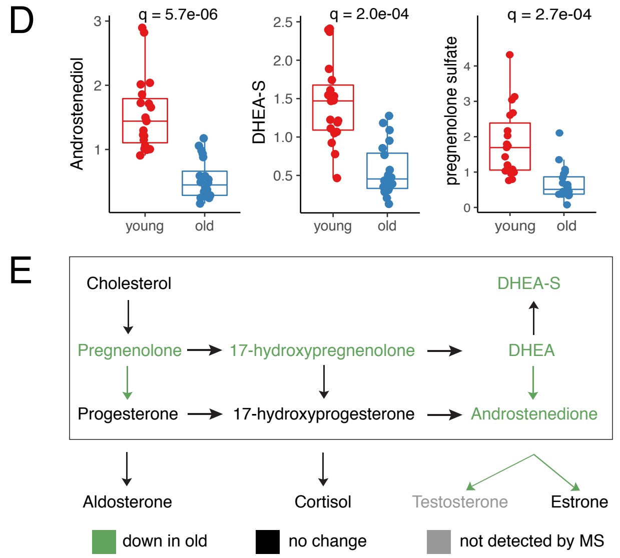


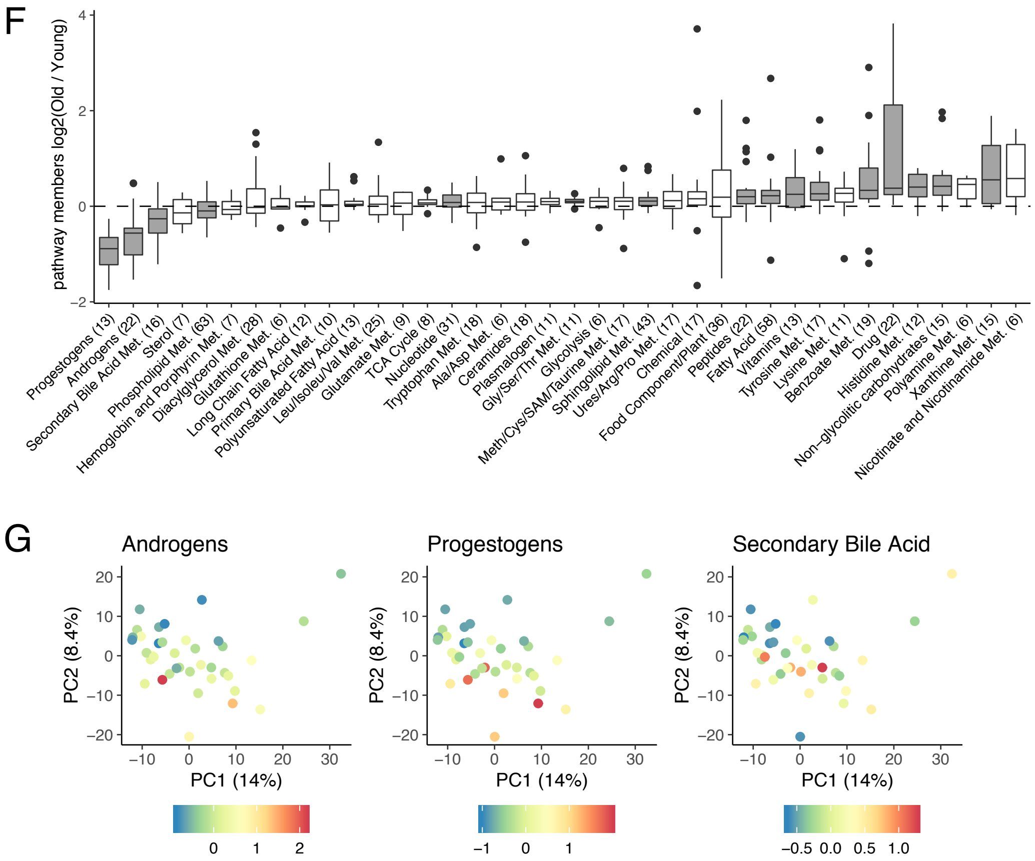

### Figure S3

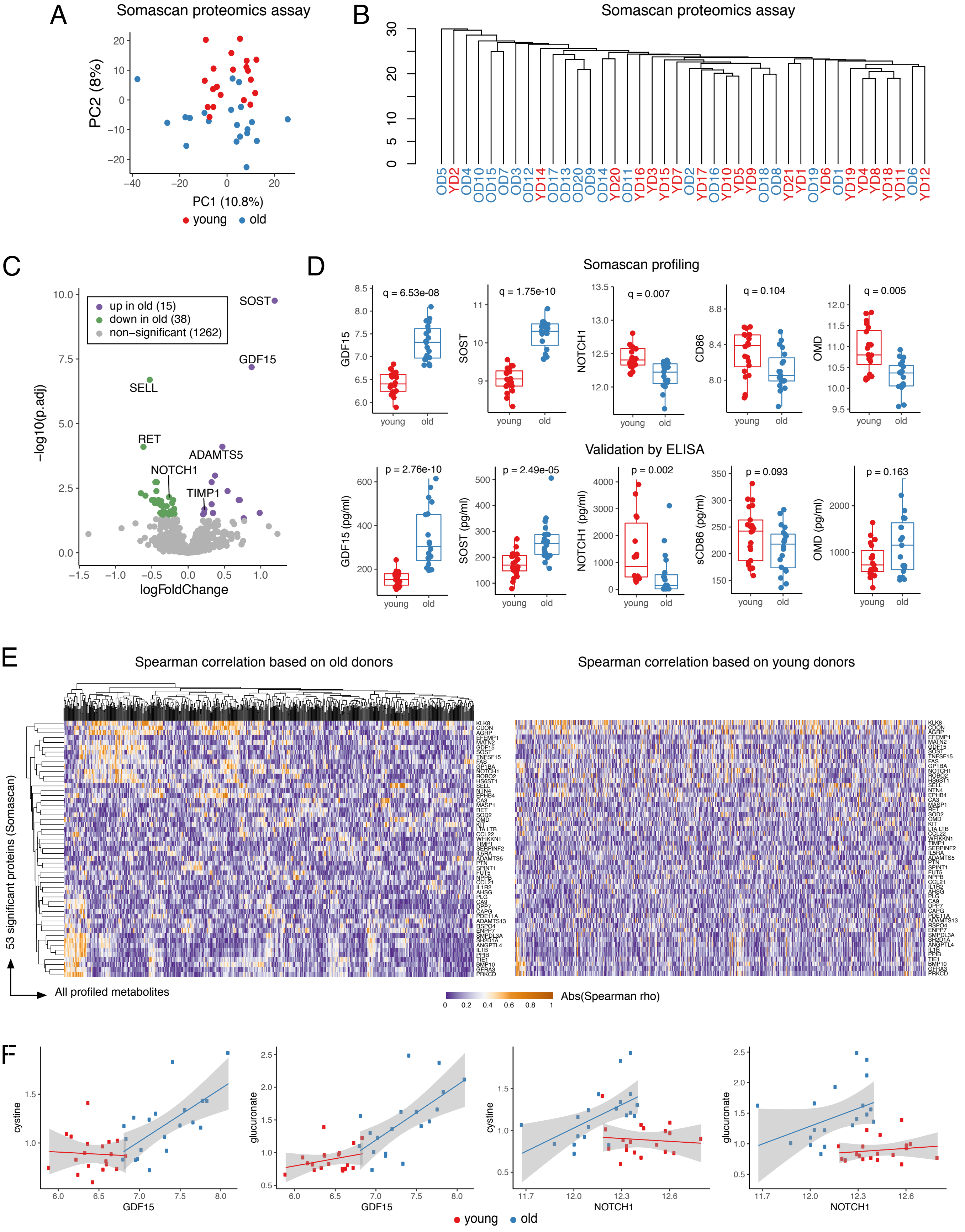
A
Somascan proteomics assay
PC2 (8%)
PC1 (10.8%)
young
old
B
Somascan proteomics assay
OD5 YD2 OD4 OD10 OD15 OD7 OD3 OD12 YD14 OD17 OD13 OD20 OD9 OD14 YD20 OD11 YD16 YD3 YD15 YD7 OD2 YD17 OD16 YD10 YD5 YD9 OD18 OD8 YD21 YD1 OD19 YD6 OD1 YD19 YD4 YD8 YD18 YD11 OD6 YD12
C
up in old (15)
down in old (38)
non−significant (1262)
−log10(p.adj)
logFoldChange
SOST
GDF15
SELL
RET
ADAMTS5
NOTCH1
TIMP1
D
Somascan profiling
q = 6.53e-08
q = 1.75e-10
q = 0.007
q = 0.104
q = 0.005
GDF15
SOST
NOTCH1
CD86
OMD
young
old
Validation by ELISA
p = 2.76e-10
p = 2.49e-05
p = 0.002
p = 0.093
p = 0.163
GDF15 (pg/ml)
SOST (pg/ml)
NOTCH1 (pg/ml)
sCD86 (pg/ml)
OMD (pg/ml)
E
Spearman correlation based on old donors
Spearman correlation based on young donors
53 significant proteins (Somascan)
All profiled metabolites
Abs(Spearman rho)
F
cystine
glucuronate
GDF15
NOTCH1
young
old

### Figure S4

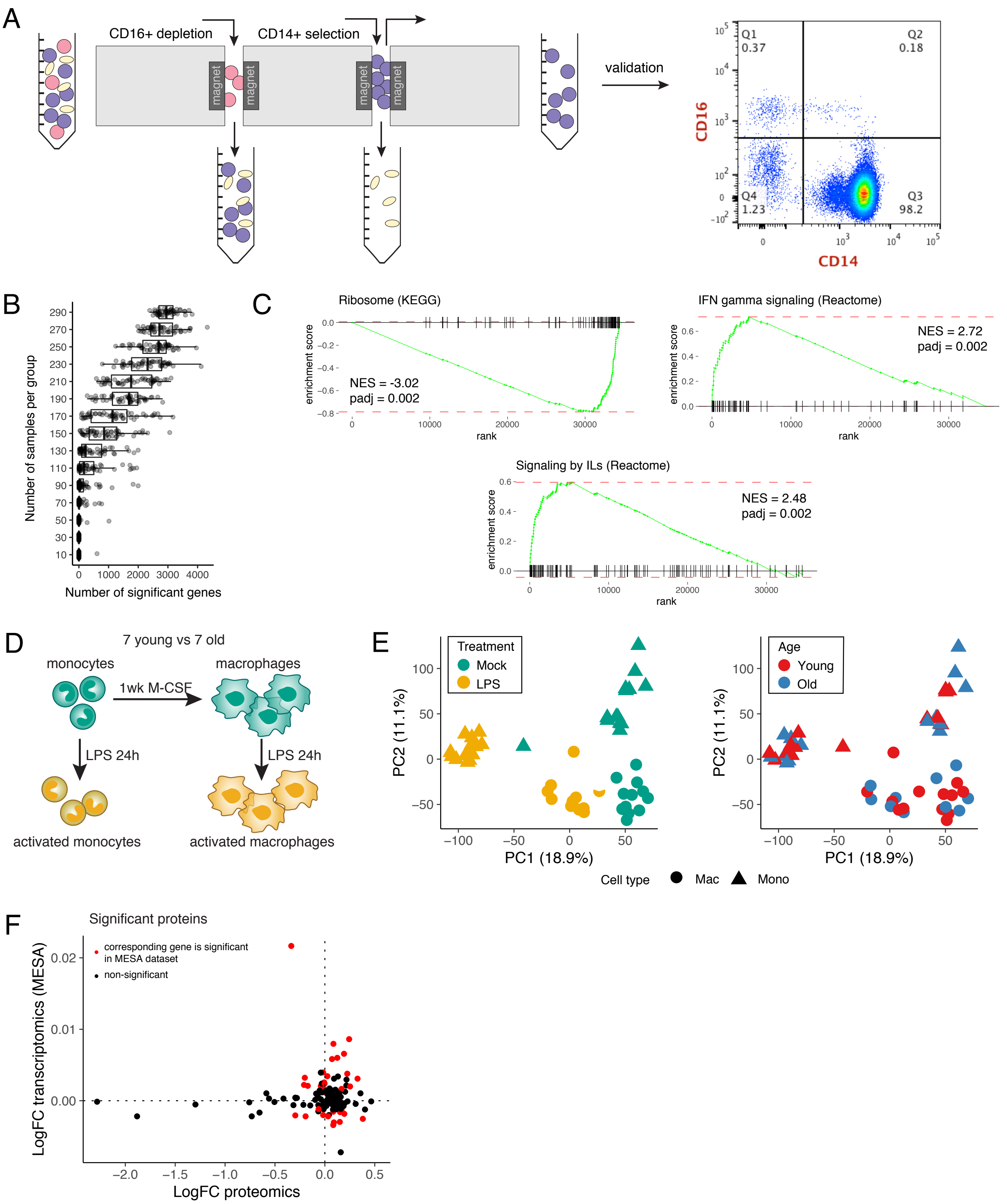
A
CD16+ depletion
CD14+ selection
magnet
validation
Q1 0.37
Q2 0.18
Q4 1.23
Q3 98.2
CD16
CD14
B
Number of samples per group
Number of significant genes
C
Ribosome (KEGG)
NES = -3.02
padj = 0.002
IFN gamma signaling (Reactome)
NES = 2.72
padj = 0.002
Signaling by ILs (Reactome)
NES = 2.48
padj = 0.002
enrichment score
rank
D
7 young vs 7 old
monocytes
macrophages
1wk M-CSF
LPS 24h
activated monocytes
activated macrophages
E
Treatment
Mock
LPS
Age
Young
Old
PC2 (11.1%)
PC1 (18.9%)
Cell type
Mac
Mono
F
Significant proteins
corresponding gene is significant in MESA dataset
non-significant
LogFC transcriptomics (MESA)
LogFC proteomics

### Figure S5

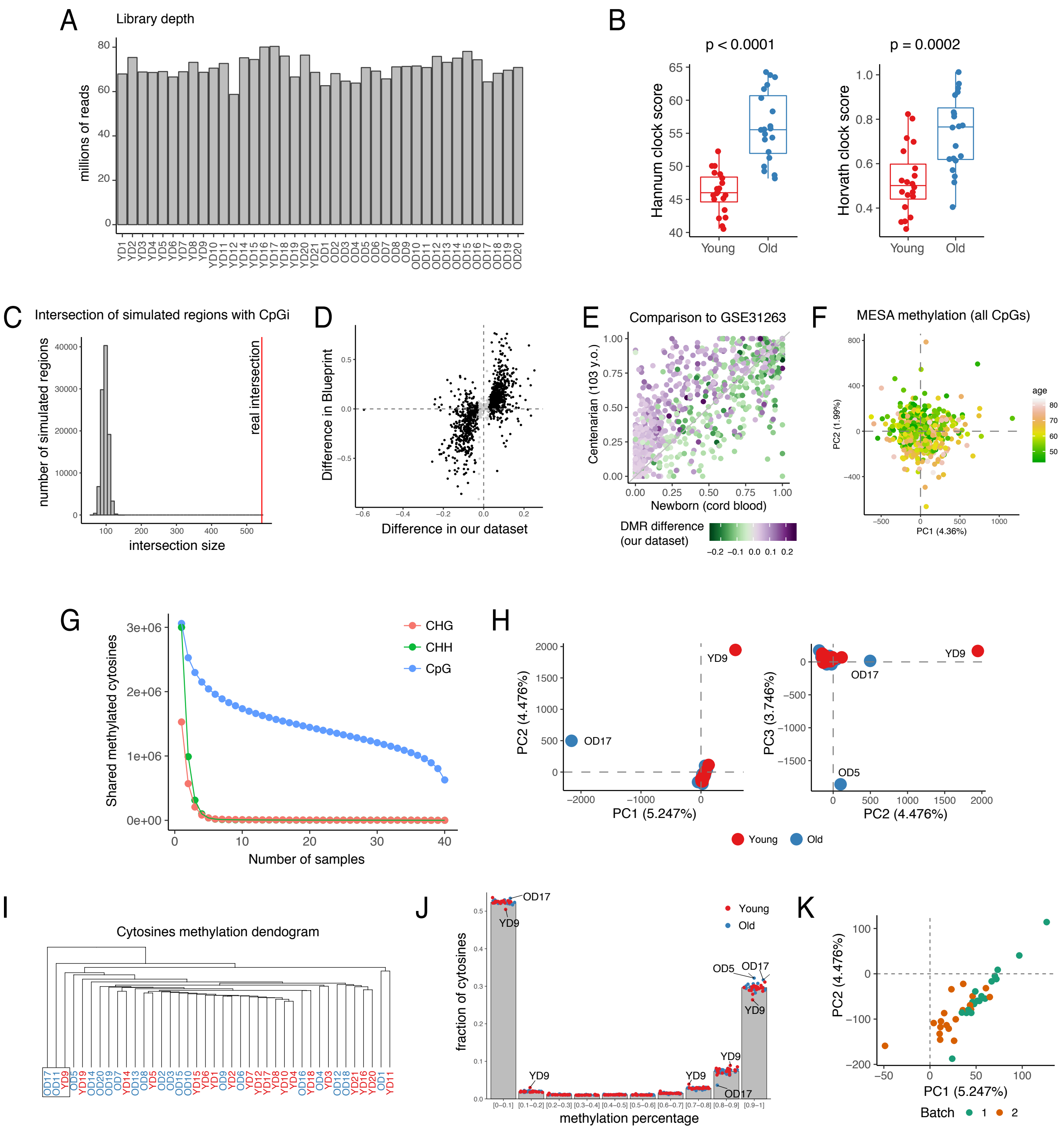

A
Library depth
millions of reads
B
p < 0.0001
p = 0.0002
Hannum clock score
Horvath clock score
Young
Old
C
Intersection of simulated regions with CpGi
number of simulated regions
real intersection
intersection size
D
Difference in Blueprint
Difference in our dataset
E
Comparison to GSE31263
Centenarian (103 y.o.)
Newborn (cord blood)
DMR difference
(our dataset)
F
MESA methylation (all CpGs)
age
PC2 (1.99%)
PC1 (4.36%)
G
CHG
CHH
CpG
Shared methylated cytosines
Number of samples
H
PC2 (4.476%)
PC1 (5.247%)
PC3 (3.746%)
PC2 (4.476%)
YD9
OD17
OD5
Young
Old
I
Cytosines methylation dendogram
J
fraction of cytosines
methylation percentage
Young
Old
K
PC2 (4.476%)
PC1 (5.247%)
Batch

### Figure S6

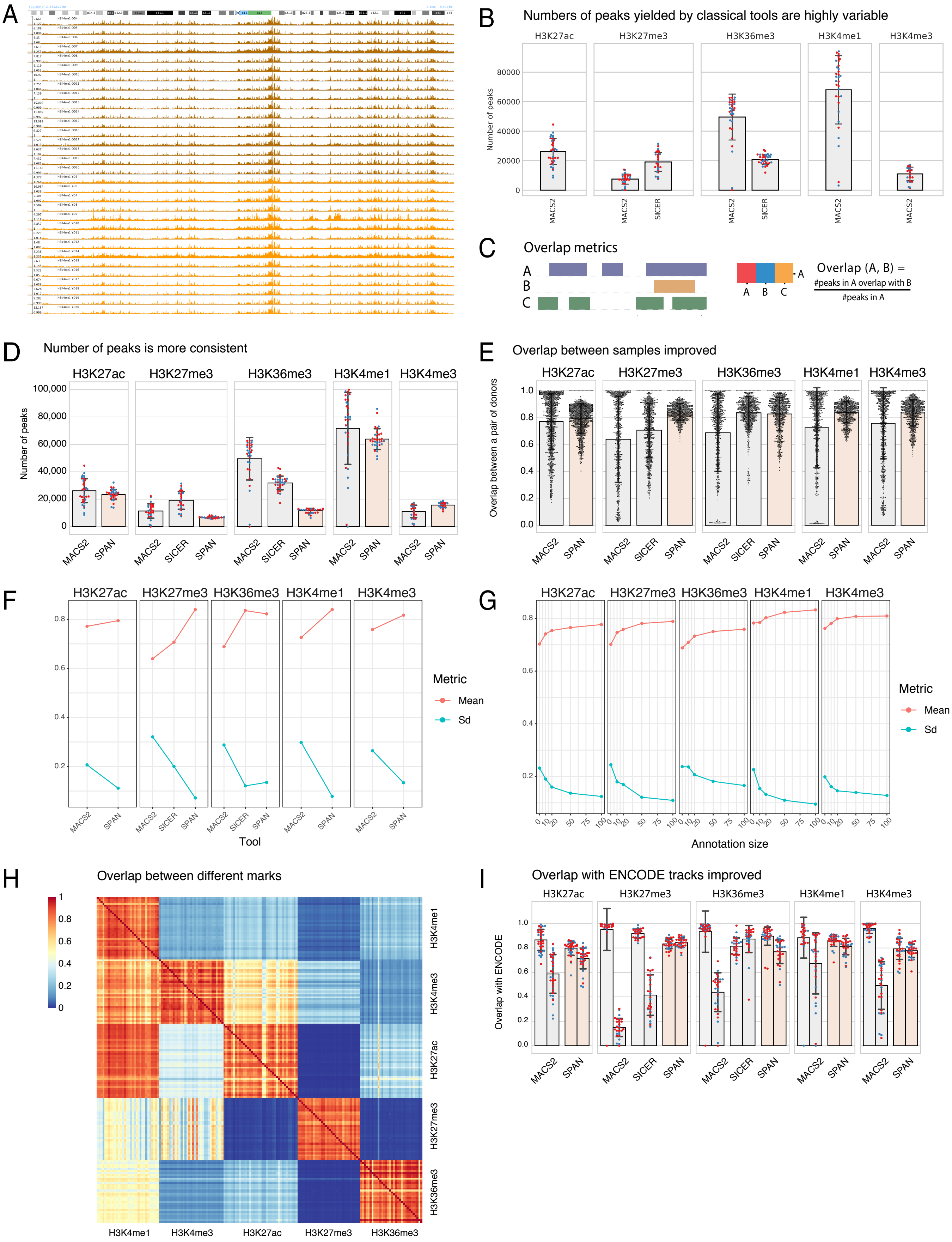

A
B
Numbers of peaks yielded by classical tools are highly variable
H3K27ac
H3K27me3
H3K36me3
H3K4me1
H3K4me3
Number of peaks
MACS2
SICER
C
Overlap metrics
Overlap (A, B) =
#peaks in A overlap with B
#peaks in A
D
Number of peaks is more consistent
SPAN
E
Overlap between samples improved
Overlap between a pair of donors
F
Metric
Mean
Sd
Tool
G
Annotation size
H
Overlap between different marks
I
Overlap with ENCODE tracks improved
Overlap with ENCODE

### Figure S7

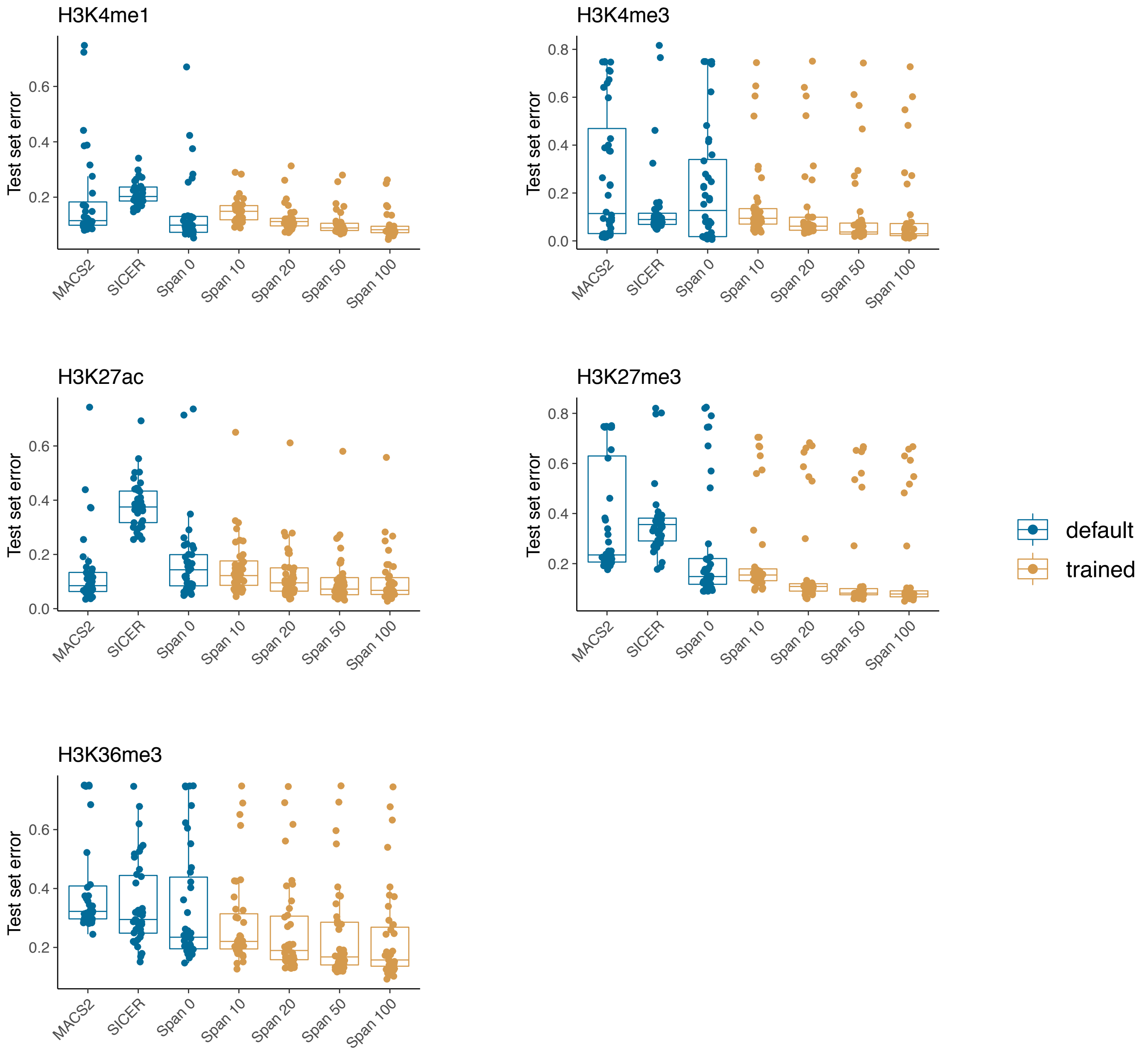
H3K4me1
H3K4me3
H3K27ac
H3K27me3
H3K36me3
Test set error
0.0
0.2
0.4
0.6
0.8
MACS2
SICER
Span 0
Span 10
Span 20
Span 50
Span 100
default
trained

### Figure S8

A

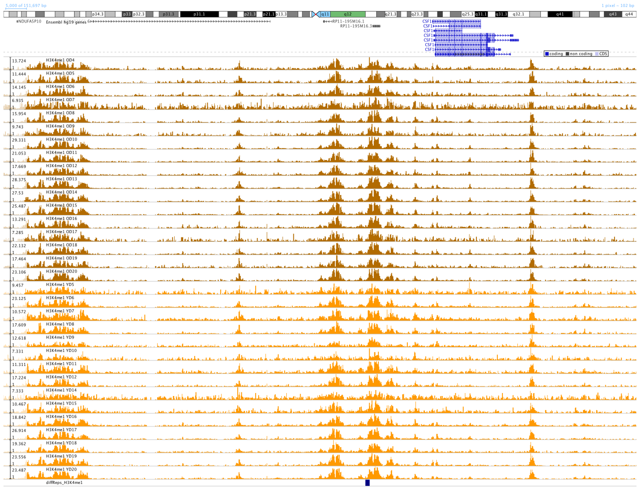

B

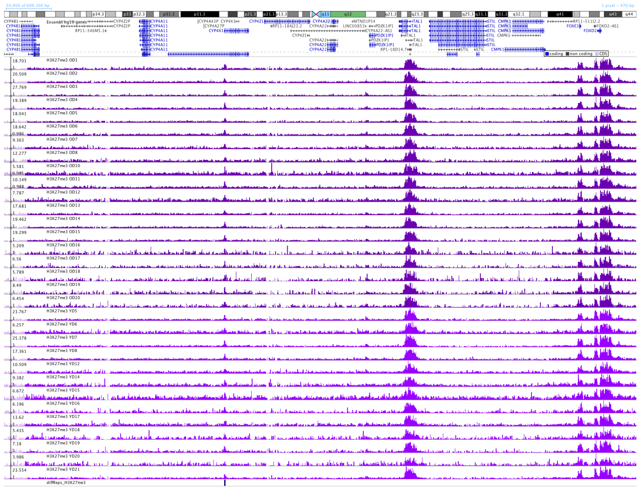

C

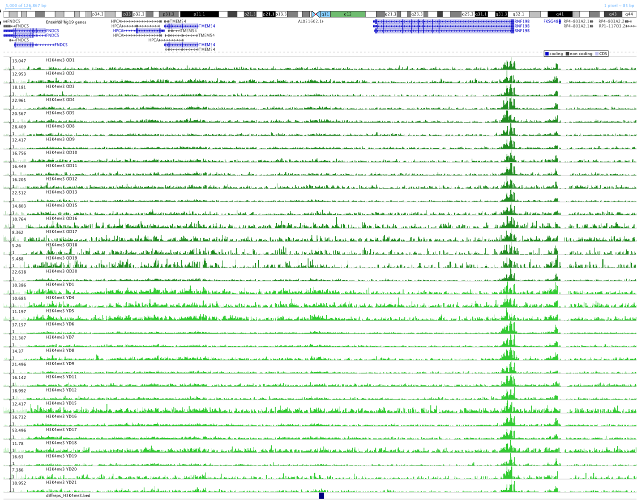

D

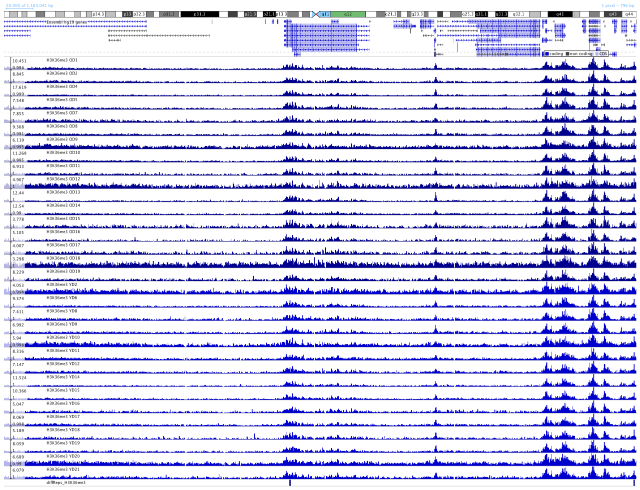

E

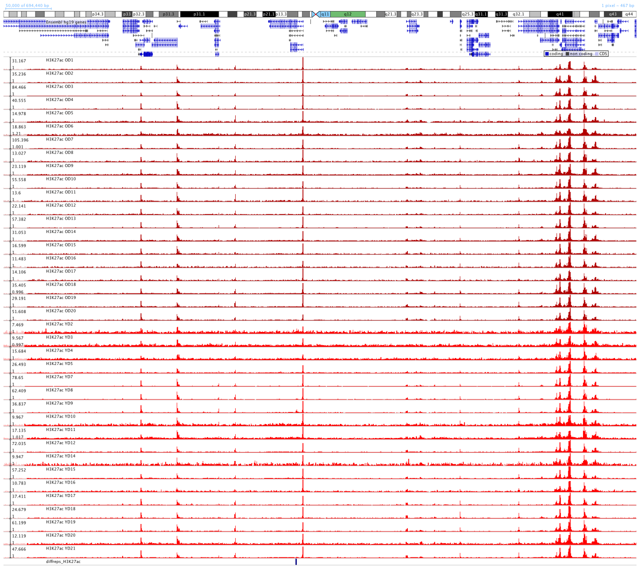

### Figure S9

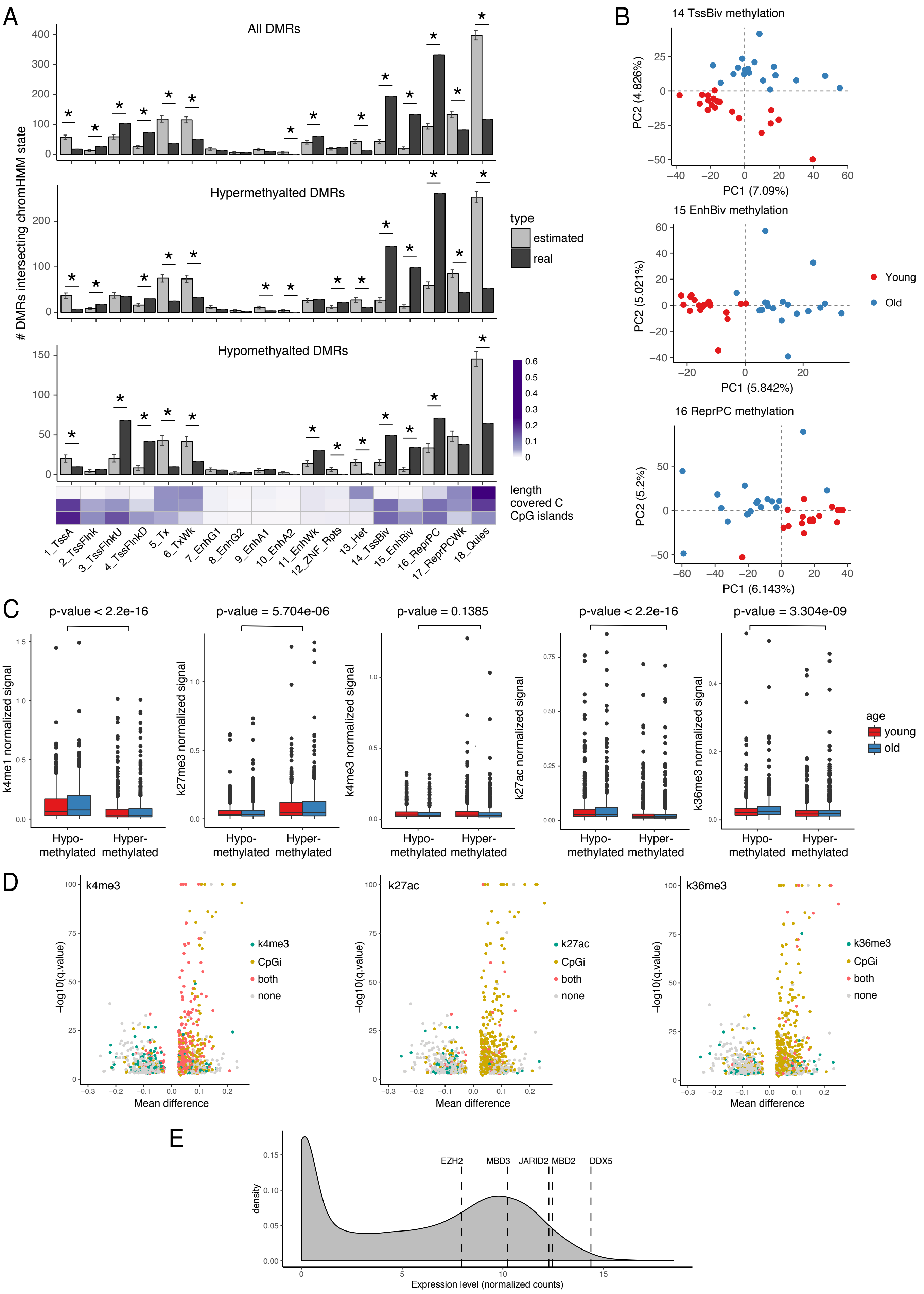
A
All DMRs
Hypermethylated DMRs
Hypomethylated DMRs
### DMRs intersecting chromHMM state
type
estimated
real
length
covered C
CpG islands
1_TssA
2_TssFlnk
3_TssFlnkU
4_TssFlnkD
5_Tx
6_TxWk
7_EnhG1
8_EnhG2
9_EnhA1
10_EnhA2
11_EnhWk
12_ZNF_Rpts
13_Het
14_TssBiv
15_EnhBiv
16_ReprPC
17_ReprPCWk
18_Quies
B
14 TssBiv methylation
PC1 (7.09%)
PC2 (4.826%)
15 EnhBiv methylation
PC1 (5.842%)
PC2 (5.021%)
16 ReprPC methylation
PC1 (6.143%)
PC2 (5.2%)
Young
Old
C
p-value < 2.2e-16
p-value = 5.704e-06
p-value = 0.1385
p-value < 2.2e-16
p-value = 3.304e-09
k4me1 normalized signal
k27me3 normalized signal
k4me3 normalized signal
k27ac normalized signal
k36me3 normalized signal
Hypo-
methylated
Hyper-
methylated
age
young
old
D
k4me3
k27ac
k36me3
−log10(q.value)
Mean difference
CpGi
both
none
E
EZH2
MBD3
JARID2
MBD2
DDX5
density
Expression level (normalized counts)

### Figure S10

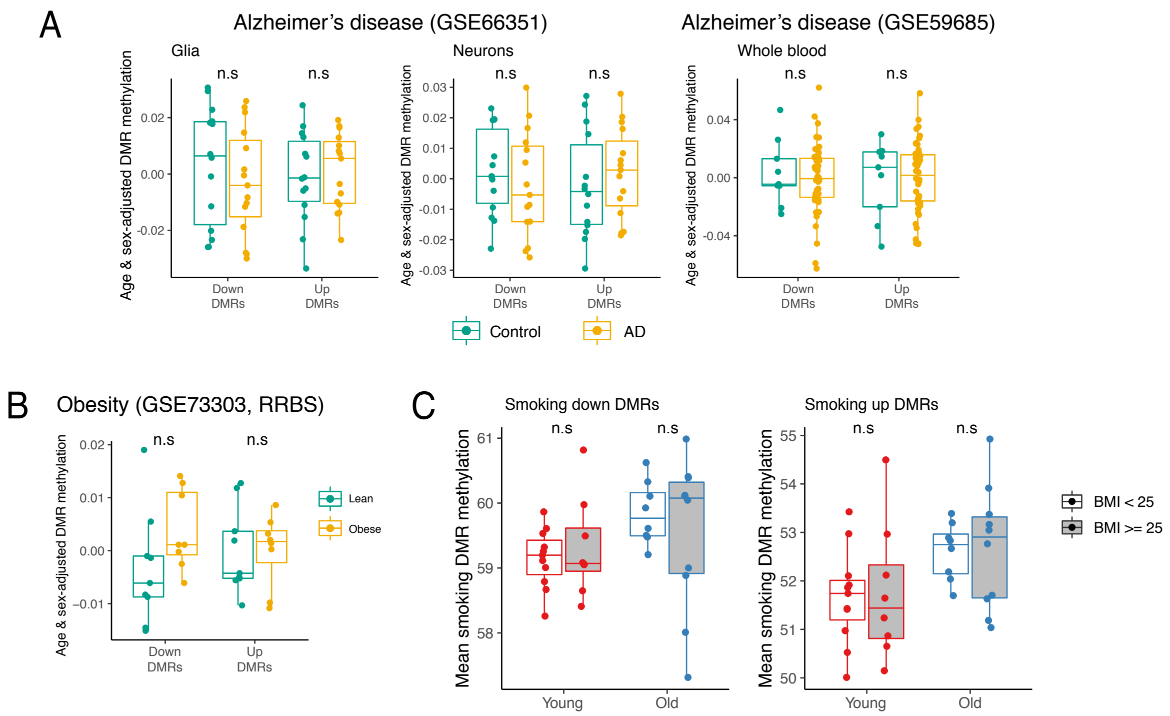
A
Alzheimer's disease (GSE66351)
Alzheimer's disease (GSE59685)
Glia
Neurons
Whole blood
n.s
Age & sex-adjusted DMR methylation
Down DMRs
Up DMRs
Control
AD
B
Obesity (GSE73303, RRBS)
Lean
Obese
C
Smoking down DMRs
Smoking up DMRs
Mean smoking DMR methylation
Young
Old
BMI < 25
BMI >= 25

### Figure S11

A

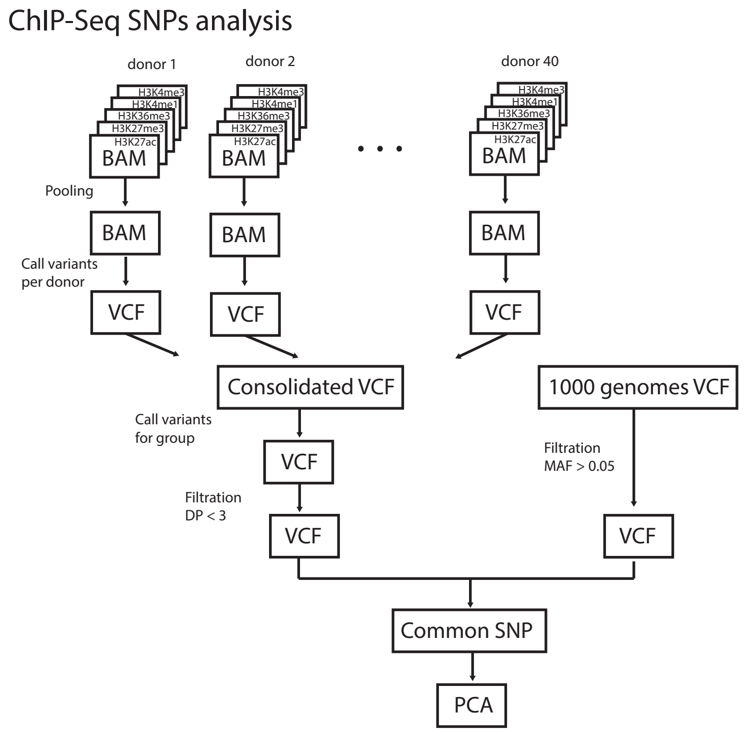


B

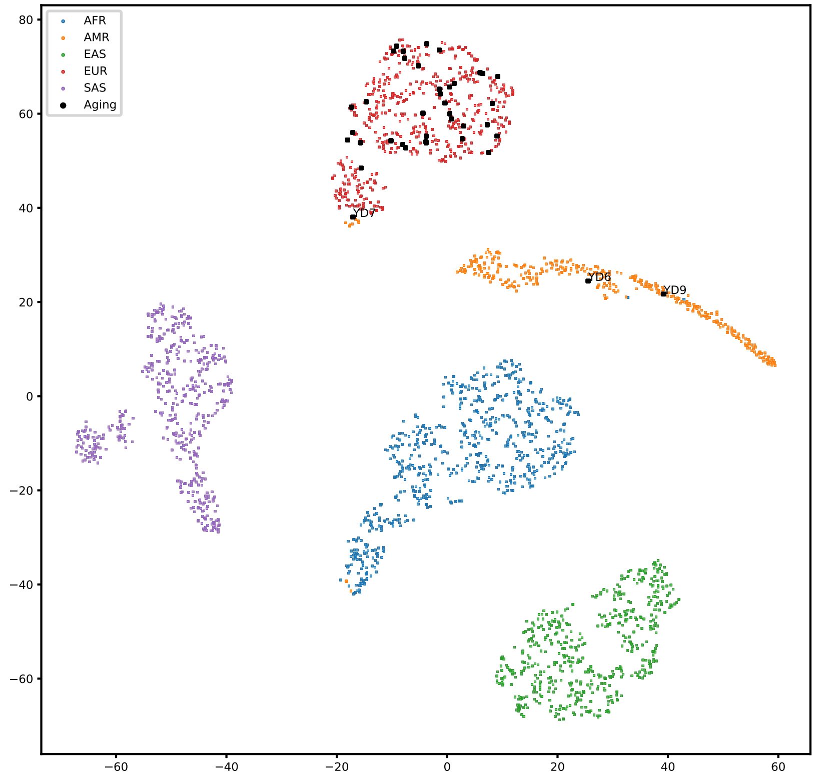
