## Supplementary materials for "Epigenetic aging of classical monocytes from healthy individuals"

**SUPPLEMENTAL METHODS**

*ChIP-Seq processing: samples overview*

Monocytes were isolated from blood samples from young (24-30 y.o.) and old (57-70 y.o.) Caucasian males as per the inclusion criteria listed in **Figure 1A** and in the methods section of the main manuscript. CD14^+^CD16^-^ monocytes were extracted from blood. We used ULI-ChIP-seq to characterize age-associated changes in the five modifications of histone 3 (H3) tails (H3K4me3, H3K4me1, H3K27ac, H3K27me3, H3K36me3) for all donors (20 young and 20 old). Overall, we generated 40 donors x 5 histone marks = 200 ChIP-seq datasets. All study participants identified themselves as Caucasians. We additionally confirmed the ancestry by comparing Single Nucleotide Polymorphisms (SNPs) from our ChIP-Seq data to SNPs commonly associated with Caucasians in 1000 Genomes dataset^1^. We pooled BAM files for all 5 modifications for each donor, called individual variants, merged them into group variants, and applied filtration DP ≥ 3. Common SNPs were used to perform Principal Component Analysis (PCA) and build t-SNE on the first five components (**Figure S11A)**. 38 put of 40 donors were identified as Europeans according to 1000 genomes data, two were mixed European American descent, **Figure S11B**.

*Establishing experimental QC pipeline (concentrations, primers, etc.)*

We used Ultra-Low-Input (ULI) ChIP-Seq protocol^2^ (<http://artyomovlab.wustl.edu/aging/methods.html>). The typical ChIP-Seq experiment requires 1-5 million cells for each run, while ULI-ChIP-Seq requires only 100k cells. However, it has certain drawbacks such as high variability of signal-to-noise ratio between samples within one library preparation. During pilot stage, we focused on optimizing ULI-ChIP-Seq protocol for isolation conditions and cell type used in our study by selecting antibody concentrations that result in the best signal to noise ratio. Final concentrations of antibodies used in our studies were: 0.05 µg for H3K27ac, 0.3 µg for H3K27me3, 0.1 µg for H3K36me3, 0.2 µg for H3K4me1 and 0.03 µg for H3K4me3.

*ChIP-qPCR Primer QC*

Using public data for CD14^+^CD16^-^ monocytes in Roadmap Epigenomics Consortium and data pilot runs we developed a collection of positive and negative control primers. These primers were used to do an additional check of signal-to-noise ratio using real-time PCR (SYBR-green) before sequencing step (see **Table S15** for sequences and locations).

*Data generation, QC and basic processing*

Initial (raw) ULI-ChIP-Seq data were subjected to standard quality control and processing steps: (1) raw reads quality, length, duplication rate, GC-content were determined for all reads; (2) alignment of raw reads on reference human genome hg19; (3) visual inspection of tracks. Reads length was 51bp, average duplication level was less than 20%, GC content was about 47%, and sequencing depth was on average ~50 million reads for all the histone modifications (**Figure 4B**). Full Reads QC data are available in the **Table S13**.

Reads quality control (step 1) revealed a problem with the first 5 base pairs (which were computationally trimmed) in all ULI-ChIP-Seq libraries, which we believe to be an artifact of the particular library preparation protocol (the same issue was observed for other cell types and species when using this protocol). Alignment on hg19 was performed using bowtie without multimappers and followed by sorting using samtools:

bowtie -p **4** -St -m **1** -v **3** --trim5 **5** --best --strata **bowtie_index** **hg19** **$**{FILE}.fastq **$**{ID}.sam

samtools view -bS **$**{ID}.sam -o **$**{ID}_not_sorted.bam
samtools sort **$**{ID}_not_sorted.bam -o **$**{BAM_NAME}

Full alignment statistics is available in **Table S13**. Aligned BAM files were analyzed by fastqc and PeakQualTools software and evaluated in accord with ENCODE data standard guidelines v4. Average alignment rate was about 85%, and most of the tracks met the standards for library depth, PCR bottlenecking coefficients, and non-redundant reads fraction. Complete code for technical ULI ChIP-Seq pipeline is available on GitHub: <https://github.com/JetBrains-Research/washu>

*Peak calling: golden standard tools*

Next, we set to perform peak calling for the compendium of ULI ChIP-Seq experiments. In 191 libraries that passed QC described above, the peaks were distinctly visible by visual inspection, yet the signal-to-noise ratio varied considerably within the cohort (see **Figure S6A**). This variability can be observed directly by applying traditional peak calling approaches such as MACS2 and SICER, which yield difference in numbers of peaks, peaks consistency, etc. We used MACS2 for all modifications and SICER for broad ones only (H3K27me and H3K36me3).

MACS2 parameters:

macs2 callpeak -t **$**{FILE} -c **$**{INPUT} -f BAM -g **hs** –broad -q **1e-4**

SICER parameters:

SICER.sh ${INPUT_FOLDER} **$**{FILE_BED} **$**{INPUT_BED} ${OUTPUT_FOLDER} **hg19** **1** **200** **150 2.7e9** **600** **1e-6**

Traditional peak calling approaches (MACS2, SICER) produced very inconsistent number of peaks for all histone marks (**Figure S6B**), which made it impossible to analyze the difference between two cohorts. We analyzed overlap between all pairs of donors and found that the average overlap rate was as low as 30% for some cases when samples were analyzed by MACS2 or SICER (**Figure S6E**), while visual data inspection suggested strong concordance between different samples. Therefore, we concluded that traditional peak calling approaches were not optimally suited for the analysis of large cohorts of ULI-ChIP-seq data.

Hocking et al ^3^ proposed a method to estimate the best parameters of peak callers using supervised labels. In this approach, user manually creates peak labels and then this information is used to find optimal peak caller parameters for all samples. The original approach was designed for conventional ChIP-Seq experiments and generated a single set of parameters for the whole group of samples. While introducing labeling information aids significantly in amplifying the consistency of peak calling, having a single set of parameters across all samples cannot be used in the context of multiple ULI-ChIP-Seq samples due to high variability of the signal-to-noise ratio between ULI-ChIP-Seq samples.

Thus, based on the concepts proposed by Hocking et al, we developed **SPAN** – a novel semi-supervised machine learning peak calling algorithm. In SPAN, we preprocess each sample separately to train the underlying statistical model. For each chromatin mark, the labeling set, that is common for all samples, is used to optimize parameters individually for each sample.

One of the major drawbacks of the semi-supervised methods is the technical difficulty of creating label annotations. A researcher has to visualize tracks in a separate genome browser, find peaks, and create text file with labeling set separately. This precludes wide-spread usage and limits the reproducibility of the data analysis. Thus, we created a dedicated genome browser, **JBR Genome Browser**, where data exploration and annotation can be processed simultaneously within one environment inside a single application.

We integrated SPAN and JBR Genome Browser to perform on-the-fly peak calling using uploaded SPAN model and either uploaded or created within the JBR Genome Browser labeling set. Such workflow allows for effortless semi-supervised peak calling and can serve as a scaffold for development of other semi-supervised approaches.

*SPAN – algorithm: overview*

The pipeline consists of 3 main steps: (1) creating SPAN model (done with SPAN in command line environment starting from bam or wig files), (2) creating manual annotation labels (can be conveniently done in JBR Genome Browser or manually), and (3) performing SPAN tuning procedure for all the ULI-ChIP-Seq tracks independently.

(1) Creating SPAN model:

For each BAM file or pair of signal and input (control) BAM files SPAN creates a 3-state HMM model that best fits the data. This part is done in the command line environment and can be a part of standardized ChIP-Seq processing routine.

(2) Creating a labeling file in BED format:

Four types of labels are used:

- *peaks*: there is at least one peak in the labeled area
- *noPeaks*: there are no peaks in the labeled area
- *peakStart*: exactly one peak **starts** in the labeled area
- *peakEnd*: exactly one peak **ends** in the labeled area

The label *peaks* is satisfied by any positive number of peaks within the label area, and it guards against too conservative calling; *noPeaks* is satisfied by no peaks intersecting the area, and it guards against too liberal calling; *peakStart* and *peakEnd* are satisfied by exactly one peak starting or ending in the area and guards against calling peaks that are too narrow.

(3) Producing final peaks optimized for signal-to-noise ratio:

BED file with labels is then passed as an input (control) file to SPAN tool whether in command line environment or within JBR Genome Browser. Given labeling BED file and model file, SPAN iterates through model parameters to find optimal parameter set that minimizes a percentage of unsatisfied labels for each ‘specific peak calling instance’. Each peak calling instance corresponds to distinct values of parameters representative of signal-to-noise ratio, peak width etc. Final error score is the ratio of number of errors to the total number of labels. SPAN iteratively varies parameter values to find the ones that optimally fit labeling pattern of the specific sample. We found that number of labels sufficient for robust tuning procedure is ~50-100.

*SPAN – algorithm: creating unsupervised model of the data*

Each chromosome is split into bins, default bin size is 200 nucleotide base pairs (other values can be passed as SPAN parameter), which loosely corresponds to the size of DNA molecule curl (loop) around histone protein complex. Unique tags coverage is calculated for each bin, producing an integer vector. Tags are shifted according to the specified fragment size. If the control file is not specified, these vectors serve as an input to the next step. If the control file is specified, the same procedure is done for control tags. The treatment and control vectors are then combined through the following formula (adapted from DiffBind):

scale = min (1, treatment_library_depth / control_library_depth)

score_i = max (0, [treatment_i - control_i * scale])

We then fit a 3-state HMM to the data from the previous step using a standard Baum-Welch method. The states are denoted NULL, LOW and HIGH. The NULL state always emits zero, while LOW and HIGH states have negative-binomially distributed emissions with parameters that are fitted. The idea here is that NULL captures the bins that have zero coverage, for example, due to being located inside a genomic repeat and thus not containing unique sequences, LOW captures the result of non-specific binding, i.e., noise, and HIGH captures the specific binding, i.e., signal. Each vector is treated as an output of the same HMM.

P(score_i=x | state_i=NULL) ~ δ(0, x)

P(score_i=x | state_i=LOW) ~ NB(x; m_LOW, f_LOW)

P(score_i=x | state_i=HIGH) ~ NB(x; m_HIGH, f_HIGH)

where state_i and score_i is the state and score of the i-th bin, δ is Kronecker's delta symbol, and NB denotes the discrete density function of a negative binomial distribution. We use a non-standard parametrization for the negative binomial distribution, namely, through the mean m and the number of failures f.

The upside here is that the log-likelihood estimator for ‘m’ is directly calculable using a method of moments. F estimator, on the other hand, is found via an iterative algorithm adapted from an article by Minka. The iterative Baum-Welch algorithm stops when the log-likelihood change becomes smaller than 0.1. It can happen, albeit very rarely, that the HIGH and LOW states get swapped during fitting so that HIGH state captures low-coverage bins and vice versa. To account for this, we employ a semantic check after the convergence and flip the states back if necessary.

The fitted model is then used to estimate posterior probabilities of each state for each bin. Assuming that the null hypothesis is that the state is not HIGH, we then calculate corresponding q-values and perform FDR control according to the provided threshold. The bins that passed the FDR control procedure are then aggregated: if two significant bins are separated by no more than GAP bins, they are merged, and all the bins in between are added to the output. The adjacent bins are joined into peaks, and the resulting peaks are saved to the provided output BED file.

*SPAN – algorithm: peak calling instances*

To distinguish the trained parameters of the underlying model and the user-supplied FDR control threshold and GAP size parameters, the latter are called meta parameters, since they are extraneous to the model itself and pertain only to the peak calling step. The meta parameters are optimized to minimize the training error of the called peaks relative to annotated labels. This is done by a simple grid search: a finite set of values is provided for each meta parameter, then the error rate is computed for every combination using the provided label file. Finally, the meta parameters combination that minimizes the error is determined, and the peaks called with that combination are returned as optimal.

By default, meta-parameters grid is as following (other values can be passed as parameters):

FDR: 1e-1, 1e-2, 1e-4, 1e-6, 1e-8, 1e-10, 1e-12;

GAP: 0, 5, 10, 20;

BIN: 200;

FRAGMENT: none, 150.

Example of command line:

java -jar span.jar --treatment control.bam --control treatment.bam --labels labels.bed --bed result.bed

In cases when several metaparameters combinations scores the same minimal error rate, we pick one with the smallest FDR and the biggest GAP. Additional testing showed that this strategy increased overall consistency, while preserving the same error rate. Importantly, even though multiple iterations are required to find optimal sets of parameters, SPAN works much faster than MACS2 and SICER in all the conditions, since it uses parallel computations for model training, both multithreading and specialized processor extensions like SSE2, AVX, etc. The pipeline was designed to reuse all the intermediate results (trained model) making optimization of meta-parameters blazing fast.

Parallel computations were performed using an open-source library for parallel matrices computations in Kotlin programming language. The source code is available on GitHub: <https://github.com/JetBrains-Research/viktor>.

*JRB-browser – technical & basic tutorials*

JBR Genome Browser is a new genome browser that brings together ability to browse and visualize genomic data, perform manual peak labeling, and run semi-supervised peak calling using SPAN. Additional important aspect of the browser is its extensibility: it is effortless for developers to create your own type of track view from scratch. Asynchronous rendering with the effective event system and tasks management makes it extremely fast. You can open up to a hundred different tracks and work with them without any significant lags. JBR Genome Browser can be used for visual peak calling; it has dedicated peak annotation mode for creating labels and peak calling. We provide a small example of the semi-supervised SPAN peak calling pipeline. The browser and all necessary test files can be found here: <https://artyomovlab.wustl.edu/aging/jbr.html>

Required files: file.bam, hg19.chrom.sizes.
(1) Train SPAN model for given BAM file:

java -jar span.jar analyze -t file.bam --cs hg19.chrom.sizes

(2) Launch the browser:

java -jar browser.jar

(3) Load BigWig visualization file and SPAN mode to browser:

File | Load BigWig file

File | Load Span Model file

(4) Turn on Peak Annotation mode using

View | Peaks Annotation Mode

(5) Create labeling set.

Select desired genomic regions and pressing a corresponding key (s for peakStart label, e for peakEnd label, p for peaks label, n for noPeaks label) or clicking the corresponding buttons.

(6) Perform final optimized peak calling

Click “tune SPAN model” to optimize peak calling for created labeling set by iterating through FDR and GAP parameters grid. Right click on labels toolbar and choose Export to *.bed to export markup.

*SPAN peak calling – application to Aging data*

All the peak calling summary information for individual donors with numbers of peaks, lengths, a fraction of reads in peaks is available in the table **Table S16**. The statistics about labels used is collected in **Table S17**. BED files with labels are available at <https://artyomovlab.wustl.edu/aging/download_data.html#download-chipseq>

### *JBR-browser for aging data*

### For data exploration convenience we prepared a set of Genome Browser sessions (for UCSC browser, IGV browser and JBR browser), available at:

<https://artyomovlab.wustl.edu/aging/explore_chipseq.html>

Each session contains the following tracks: (1) median consensus, i.e. overlapping peaks, existing in at least half of all samples, (2) weak consensus – overlapping peaks confirmed by at least 2 samples, (3) section of Bigwigs with raw reads for young and old donors, low quality failed tracks are colored in gray, (4) section of corresponding peaks produced by pipeline, (5) ENCODE track, (6) peaks for ENCODE track and (7) – labels used for semi-supervised peak calling. IGV sessions can be opened in JBR Genome Browser as well to make it easier to modify labeling (7) and redo peak calling after downloading preprocessed models.

*SPAN study case 1: public ULI ChipSeq data (GSE63523)*

As a benchmarking, we also applied SPAN peak calling approach to publicly available ULI ChIP-Seq datasets. In the paper Chen C et al. presented an ultra-low-input micrococcal nuclease-based native ChIP (ULI-NChIP) and sequencing method, to generate genome-wide histone mark profiles with high resolution and reproducibility from as few as one thousand cells. They created data for both broad and narrow histone marks from 10^3-10^6 cells. Among others in GSE63523, there are H3K4me3 tracks for 100, 10 and 5 thousand cells and H3K27me3 for 100, 10 and 1 thousand cells. We used these tracks to evaluate the semi-supervised approach in extreme conditions.

SPAN pipeline yielded ~10 thousand H3K3me3 peaks and ~20 thousand H3K27me3 peaks for 100 thousand cells, which is comparable with ENCODE and aging study peaks. Also, we used MACS2 with default settings FDR=0.05 and FDR=1e-6.

SPAN produced approximately the same number of peaks for H3K27me3 for 100 and 10 thousand cells (~20000) and half of the peaks for 1 thousand cells, while MACS2 with relaxed FDR produces too many peaks in all cases and too small number of peaks for 10 thousand (~7000). The situation is even worse for H3K4me3 modification, MACS2 produced too many peaks for relaxed and too less for stringent FDR control, while SPAN produced ~17000 peaks, which is comparable to what we see in other study cases and aging project data. All the peak callers failed given smaller than 100 thousand cells for H3K4me3, which suggests that this is a minimum required for high-quality peak calling.

Labels, reads, and peaks visualization are available in the sessions at

<https://artyomovlab.wustl.edu/aging/study_cases.html#span-usage-uli>

*SPAN study case 2: public McGill dataset*

SPAN tuning mode was inspired by the article by T. D. Hocking et al. Authors investigated the benefits of tuning approach for golden standard peak callers. Since the authors were affiliated with McGill University (Montréal, Canada) and used the McGill Epigenomics Mapping Centre data, we will further refer to this research as “the McGill experiment.”

The Hocking et al. experimental data consisted of 37 human cell samples. T cells, B cells, and monocytes were grouped as "immune" (27 samples), and kidney, skeletal muscle, and leukemic B cells were grouped as "other" (10 samples). Each sample was subjected to H3K4me3-specific ChIP-seq procedure, and 29 samples were additionally subjected to the H3K36me3-specific procedure (21 immune and eight other).

Each of the four participating researchers then created one or several datasets, comprising all tracks for a single histone modification (H3K4me3 or H3K36me3) and a single cell group (immune or other). A set of cell-type specific labels was created for each dataset. Each label belongs to one of the same four types that we use for tuning SPAN (peaks, noPeaks, peakStart, peakEnd). For example, the H3K36me3_AM_immune dataset consists of 83 labels created by a researcher codenamed AM for the 21 H3K36me3 tracks created from immune cell samples. In total, seven datasets were generated.

Authors selected several peak callers and fixed a parameter grid for each of them. They then called the peaks for each ChIP-seq track of each dataset for each parameter value of each caller. Our proposed tuning method somewhat differs from that employed by Hocking et al. Namely, we tune each track individually, meaning we assign an individual optimal meta parameter value to each track in each dataset, while the McGill experiment selects single meta parameter that optimizes the total dataset error. Our approach offers tangible benefits for SPAN since it has been empirically confirmed that different quality tracks require different FDR cutoffs. Other peak callers seemed to benefit from this as well; the optimal parameter value varied widely from track to track for all the golden standard peak callers, and the total error was significantly improved by individual tuning, see **Table S18**.

The IGV and UCSC sessions for the relevant data can be downloaded at

<https://artyomovlab.wustl.edu/aging/study_cases.html#span-usage-mcgill>

*Differential ChIP-Seq*

The section below briefly describes parameters used for all the differential ChIP-Seq tools:

DiffBind requires predefined peaks for every donor. We used peaks previously called by SPAN pipeline and parameters:

score = DBA_SCORE_TMM_MINUS_FULL

fragmentSize = 125

ChIPDiff doesn’t support biological replicates so pooled samples were used as input. BAM was converted to TAG file using fragmentSize = 125. Default parameters were used:

maxIterationNum   500

minP              0.95

maxTrainingSeqNum 10000

minFoldChange     3

minRegionDist     1000

ChIPDiff Y_tags.tag O_tags.tag hg19.chrom.sizes config.txt ${NAME}

MACS2 bdgdiff used BedGraph files produced by MACS2 on samples polled by cohort.

macs2 bdgdiff \

   --t1 Y_treat_pileup.bdg --c1 Y_control_lambda.bdg \

   --t2 O_treat_pileup.bdg --c2 O_control_lambda.bdg \

   --d1 ${CONTROL_Y} --d2 ${CONTROL_O} --o-prefix ${NAME}

DiffReps has different regimes for broad and narrow peaks, we used pooled BAM converted to BED.

Parameters used for broad peaks:

diffReps.pl \

      -co YD*.bed --bco YD_input.bed \

      -tr OD*.bed --btr OD_input.bed \

      --chrlen hg19.chrom.sizes \

      -re diff.nb.txt \

      --mode block --nsd broad \

      --nproc 8

For narrow peaks we used parameters:

diffReps.pl \

      -co YD*.bed --bco YD_input.bed \

      -tr OD*.bed --btr OD_input.bed \

      --chrlen hg19.chrom.sizes \

      -re diff.nb.txt \

      -me nb \

      --nproc 8

We explored the output of differential ChIP-Seq peak calling visually and were not able to detect any significant changes in chromatin modifications between two cohorts. The lack of the difference was confirmed by all of the peak callers used except diffReps. Visual analysis of top significant diffReps peaks suggests that those found are method artifacts rather than real difference.

Following images show regions with top p-value differential peaks reported by diffReps **(Figure S8)**:

H3K27ac chr13:50204201-50205200

H3K27me3 chr10:102098001-102108000

H3K4me1 chr10:103597001-103607000

H3K4me3 chr1:33391201-33392200

H3K36me3 chr10:126385001-126396000

**SUPPLEMENTARY FIGURE LEGENDS**

**Figure S1: (A)** Blood cytokine levels measured by bioplex assay and **(B)** Blood differentials obtained using Hemavet. Normal ranges for humans are shown below the boxplots. P-values for all comparisons were calculated using two-sided Mann-Whitney U test.

**Figure S2: (A)** PCA of standardized levels of metabolites in plasma. Each dot represents one donor, percentage of variance explained by principal components is shown in brackets. **(B)** Dendrogram produced by unsupervised hierarchical clustering of metabolic data. Clustering using average algorithm and Euclidian distance as the distance metric. **(C)** List of significantly different metabolites. **(D)** Q-values of selected differentially regulated metabolites from sex steroids synthesis pathway (two-sided Mann-Whitney U test and Benjamini–Hochberg correction for multiple testing). **(E)** Schema of sex steroids synthesis. Green font indicates significant decrease of corresponding metabolite in old group. Green arrow depicts intermediate pathway metabolites that are significantly decreased. Metabolites that were not detected by LC-MS profiling are shown in grey. **(F)** Pathway analysis of metabolic data. Each boxplot summarizes log2FC of all members of the corresponding pathway. Grey color indicates pathways with mean log2FC significantly different from zero (two-sided Mann-Whitney U test and Benjamini–Hochberg correction for multiple testing). **(G)** PCA as in **Figure S2A**. Each dot represents one sample. Z-scores were calculated for all metabolites. For each sample, color of the dot represents averaged z-scores of all metabolites belonging to the pathway.

**Figure S3: (A)** PCA of standardized proteins levels in plasma (Somascan). Each dot (data point) represents one donor, and percentage of variance explained by principal components is shown in brackets on each axis. **(B)** Dendrogram produced by unsupervised hierarchical clustering of Somascan data. Clustering using average algorithm and Euclidian distance as the distance metric. **(C)** Differential analysis results for plasma proteomic profile: volcano plot. Each dot represents a single protein. Significantly different proteins are highlighted. Adjusted p-values and logFC were calculated by Limma package. **(D)** ELISA validation of selected proteins. Q-values for Somascan results (Limma package and Benjamini–Hochberg correction for multiple testing) are shown in the top panel and ELISA validation of the same targets are shown below. P-values were calculated using two-sided Mann-Whitney U test. **(E)** Heatmap representation of absolute values of Spearman’s correlation coefficients (rho) between plasma proteins (rows) and plasma metabolites (columns) calculated within old (left panel) and young (right panel) cohorts. Clustering of old cohort rows and columns was done using complete algorithm and Euclidian distance as a metric. Order of rows and columns in young cohort heatmap matches order established for the old cohort. **(F)** Each point represents one donor. Smoothing was done by *lm* function separately in young and old groups.

**Figure S4: (A)** Left panel: schematic representation of CD14^+^CD16^-^ monocytes isolation using magnetic beads. Right panel: flow cytometry validation and estimation of purity. **(B)** Number of significant genes detected after downsampling MESA dataset (youngest and oldest 25%). Downsampling was repeated 50 times for each group size. **(C)** GSEA enrichment curves illustrate pathways that significantly change with age in MESA dataset. **(D)** Monocytes were differentiated into macrophages by one-week incubation with M-CSF. Both cell types were stimulated by LPS for 24 hours. **(E)** PCA of normalized expression levels estimated by RNA-seq for monocyte differentiation and activation experiment. Each dot represents one sample, percentage of variance explained by principal components is shown in brackets. Dots are colored by treatment (left panel) or donor age (right panel). Dot shape corresponds to cell type. **(F)** Each dot represents a monocyte protein significantly different between age groups. LogFC proteomics (x axis) as in **Figure 2F**, LogFC transcriptomics as in **Figure 2C**.

**Figure S5: (A)** Library depth for each sample. **(B)** Hannum and Horvath methylation clocks for old and young groups. Methylation levels of CpGs that were used in the model but were not covered in our eRRBS data were imputed using mean methylation of [-100kb; +100kb] region around the CpG. CpG methylation was set to zero if imputation was not possible. P-values were calculated using two-sided Mann-Whitney U test. **(C)** Enrichment of DMRs in CpG islands. Histogram shows distribution of simulated intersection sizes (100,000 random simulations), red line – real intersection. **(D)** Comparison to Blueprint dataset. Right panel: each dot represents one DMR detected in our dataset. X axis – difference between old and young cohorts in our dataset, Y axis – difference between old donors and cord blood from Blueprint. **(E)** Plot as in right panel of **Figure 3G** for a newborn vs centenarian WGBS dataset (GSE31263). **(F)** PCA on MESA data as in **Figure 3I** using all cytosines profiled by DNA methylation array. Color represents donor age. **(G)** Number of methylated (methylation level > 0) cytosines in CpG, CHG and CHH context shared by one to 40 samples. **(H)** PCA of CpGs methylation levels from old (blue) and young (red) groups. Each dot represents one sample, percentage of variance explained by principal components is shown in brackets. **(I)** Dendrogram produced by unsupervised hierarchical clustering of the samples. Each sample described as a vector of CpG methylation levels. Clustering using Ward algorithm and Manhattan distance as the distance metric. Outliers (OD11, OD17, YD9) are labelled. **(J)** Distribution of CpG methylation levels. For each segment fractions of CpGs with corresponding methylation level in each sample are shown by dots. Bar shows average fraction of CpG across all samples. Outlying samples are labelled. **(K)** PCA of CpGs methylation levels. Outliers (OD11, OD17, YD9) were excluded. Each dot represents one sample, samples from the first batch are green, from the second batch are orange. Percentage of variance explained by principal components is shown in brackets.

**Figure S6: (A)** Snapshot of the H3K4me1 tracks across all donors shows distinct signal for all samples with visible variability in the signal-to-noise ratio. **(B)** Number of peaks for each mark in each donor yielded by classical peak calling tools. **(C)** Schematic representation of overlap metric used in **Figures S6E** and **S6H**. **(D)** Number of peaks yielded by SPAN, MACS2 and SICER. **(E)** Overlap between all pairs of samples for peaks generated by SPAN, MACS2 and SICER. SICER was used for wide modifications only. **(F)** Summary for **Figure S6E**. Mean and standard deviation (Sd) of overlaps between samples are shown. **(G)** Overlap characteristics as in **Figure S6F** for SPAN runs with various annotation sizes. **(H)** Directional overlap of SPAN peaks between all samples and all histone modifications. **(I)** Overlap with ENCODE CD14^+^ monocytes data for different peak calling approaches.

**Figure S7**: Test set errors for golden standard tools (MACS2, SICER) as well as SPAN trained with various numbers of labels.

**Figure S8:** Visualization peaks reported by diffReps as differential between young and old cohorts (peaks with smallest p-values selected). **(A)** H3K4me1 **(B)** H3K27me3 **(C)** H3K4me3 **(D)** H3K36me3 **(E)** H3K27ac.

**Figure S9: (A)** Enrichment of DMRs in chromatin state segments from ENCODE ChromHMM partition. Upper: for each chromatin state dark grey bar represents number of DMRs intersecting at least one segment of the state. Light grey bar shows expected number of intersections estimated by 100,000 random simulations. Error bars show SD. Results shown for all DMRs, hypermethylated DMRs only and hypomethylated DMRs only. Chromatin states that are significantly over- or under-represented among DMRs are marked by asterix (*). Bottom: heatmap shows general statistics for each chromatin state. Values are normalized within each row. **(B)** PCA of standardized methylation levels in old (blue) and young (red) groups for three chromatin states: bivalent Tss (14 TssBiv), bivalent enhancers (15 EnhBiv) and Polycomb-repressed regions (16 ReprPC). Each dot represents one sample, percentage of variance explained by principal components is shown in brackets. **(C)** Intensity of H3K4me1, H3K27me3, H3K4me3, H3K27ac, and H3K36me3 signals was calculated for each DMR via Diffbind package, normalized with respect to the DMR length and averaged across the cohorts. Normalized signals were compared between hypo- and hypermethylated DMRs using two-sided Mann-Whitney U test. **(D)** Volcano plot as in **Figures 4I** and **4K**, colored in accordance with intersection with H3K4me3, H3K27ac and H3K36me3. **(E)** Density plot protein coding gene expression with transcription factors of interest highlighted.

**Figure S10: (A)** Comparison of age- and sex-adjusted DMR mean methylation between Alzheimer’s patients and healthy controls. Each dot represents one donor. P-values calculated using two-sided Mann-Whitney U test. **(B)** Plot as in **S10A** comparing data from lean and obese donors. **(C)** Plot as in **Figure 6G**. Mean methylation of smoking DMRs in our donors split by age and BMI. P-value was calculated using two-sided Mann-Whitney U test.

**Figure S11: (A)** Scheme of SNP calling analysis pipeline **(B)** tSNE plot for samples from the 1000 genome data and donors participating in this study (black dots)

**SUPPLEMENTARY TABLE LEGENDS**

**Table S1:** Basic donor information and blood differential (cell counts, Hb and HCT levels)

**Table S2:** Cytokine bioplex assay data for all donors (IFN-γ, IL-10, IL-12p70, IL-13, IL-1β, IL-2, IL-4, IL-6, IL-8, TNF-α).

**Table S3: (A)** Scaled intensities of the metabolites in the plasma for all donors (734 metabolites). **(B)** Differential analysis results.

**Table S4:** SomaScan proteomics array data for all donors: **(A)** scaled data, **(B)** differential comparison statistics.

**Table S5:** ELISA validation of the most differentially present plasma proteins (sCD86, GDF-15, SOST, OMD, Notch 1).

**Table S6:** Counts for monocyte RNA-seq data. **(A)** Raw counts, **(B)** DESeq2 normalized counts, **(C)** Log2-quantile normalized counts.

**Table S7:** Monocyte RNA-seq data. DESeq2 differential analysis results between old and young cohorts.

**Table S8:** Transcriptomic data for MESA cohort. Limma differential analysis results (*gene ~ age + chip + race-gender-site* design).

**Table S9:** Monocyte proteomics data generated and analyzed by Biognosys. **(A)** Sample table, **(B)** Spectral library protein inventory, **(C)** Spectral library peptide inventory, **(D)** Differential analysis results, **(E)** Protein intensities, **(F)** Peptide intensities.

**Table S10:** RNA-seq data for monocyte differentiation and stimulation experiment (see **Figure S4D**). **(A)** Raw counts, **(B)** DESeq2 normalized counts, **(C)** Log2-quantile normalized counts.

**Table S11:** Basic QC metrics of RRBS libraries (N covered (>= 10 reads) CpG, mean CpG coverage, % mapped reads, conversion rate, library depth).

**Table S12:** Differentially methylated regions as obtained from RRBS data: **(A)** unfiltered DMRs, **(B)** Confident DMRs filtered based on ncyto ≥ 3 and abs(avdiff) ≥ 0.025.

**Table S13:** Basic characteristics and QC of the ULI-ChipSeq libraries: **(A)** FastqC output, **(B)** QC of bam files based on ENCODE standards, **(C)** alignment statistics, **(D)** QC results reported by phantompeaks.

**Table S14:** Transcription factors binding sites overrepresentation analysis: **(A)** Results for up DMRs corrected for enrichment in CpG islands, **(B)** Results for down DMRs corrected for enrichment in H3K4me1 marked regions.

**Table S15:** Positive and negative primers used for the ChIP-qPCR quality control.

**Table S16:** Peak calling summary for all samples (MACS2, SICER and SPAN).

**Table S17:** Labels used for the semi-supervised peak calling: **(A)** overall label sets statistics. Specific labels for **(B)** H3K27ac, **(C)** H3K27me3, **(D)** H3K36me3, **(E)** H3K4me1, **(F)** H3K4me3.

**Tables S18:** Comparisons of errors produced by SICER, MACS2 and SPAN on the datasets annotated by McGill team from Hocking et al.

**SUPPLEMENTARY MATERIALS REFERENCES**

1.	Genomes Project, C.*, et al.* A global reference for human genetic variation. *Nature* **526**, 68-74 (2015).

2.	Brind'Amour, J.*, et al.* An ultra-low-input native ChIP-seq protocol for genome-wide profiling of rare cell populations. *Nat Commun* **6**, 6033 (2015).

3.	Hocking, T.D.*, et al.* Optimizing ChIP-seq peak detectors using visual labels and supervised machine learning. *Bioinformatics* **33**, 491-499 (2017).
